# Structural insights into ion conduction by novel cation channel, TMEM87A, in Golgi apparatus

**DOI:** 10.1101/2023.01.03.522544

**Authors:** Ah-reum Han, Aihua Zhang, Hyunji Kang, Miguel A. Maria-Solano, Jimin Yang, C. Justin Lee, Sun Choi, Ho Min Kim

**Affiliations:** Center for Biomolecular and Cellular Structure, Life Science Cluster, Institute for Basic Science (IBS), Daejeon 34126, Republic of Korea; Global AI Drug Discovery Center, College of Pharmacy and Graduate School of Pharmaceutical Science, Ewha Womans University, Seoul 03760, Republic of Korea; Center for Cognition and Sociality, Life Science Cluster, Institute for Basic Science (IBS), Daejeon 34126, Republic of Korea; Graduate School of Medical Science and Engineering, Korea Advanced Institute of Science and Technology (KAIST), Daejeon 34141, Republic of Korea

**Keywords:** TMEM87A, Cryo-EM, Cation Channel, Golgi apparatus

## Abstract

TMEM87 family is evolutionarily conserved eukaryotic transmembrane proteins residing in the Golgi^1^. TMEM87 members play a role in retrograde transport in Golgi and are also proposed mechanosensitive ion channel implicated in cancer and heart disease^2–7^. In an accompanying study, TMEM87A is described as a voltage-gated, pH-sensitive, non-selective cation channel whose genetic ablation in mice disrupts Golgi morphology, alters glycosylation and protein trafficking, and impairs hippocampal memory. Despite the pivotal functions of TMEM87s in Golgi, underlying molecular mechanisms of channel gating and ion conduction have remained unknown. Here, we present a high-resolution cryo-electron microscopy structure of human TMEM87A (hTMEM87A). Compared with typical ion channels, the architecture of hTMEM87A is unique: a monomeric cation channel consisting of a globular extracellular/luminal domain and a seven-transmembrane domain (TMD) with close structural homology to channelrhodopsin. The central cavity within TMD is occupied by endogenous phosphatidylethanolamine, which seals a lateral gap between two TMs exposed to the lipid bilayer. By combining electrophysiology and molecular dynamics analysis, we identify a funnel-shaped electro-negative luminal vestibule that effectively attracts cations, and phosphatidylethanolamine occludes ion conduction. Our findings suggest that a conformational switch of highly conserved positively-charged residues on TM3 and displacement of phosphatidylethanolamine are opening mechanisms for hTMEM87A, providing an unprecedented insight into the molecular basis for voltage-gated ion conduction in Golgi.

## Main Text

As an integral component of the endomembrane system, the Golgi apparatus forms a semi-circle shaped series of flat sacs; cisternae, within which lipids and proteins are continuously modified and transported to meet cellular needs. To execute these functions, maintaining ion and pH homeostasis of the Golgi lumen is critical^8–10^. Hence, dysregulation of Golgi ion and pH homeostasis can alter protein trafficking and glycosylation and even lead to Golgi disorganization, a cellular phenotype associated with various diseases such as cancer and neurodegenerative disease^11–14^. While several anion channels in the Golgi membrane, including Golgi pH regulator (GPHR), mid-1-related chloride channel (MClC), and voltage-gated chloride channels (ClC-3B) are known to counterbalance for Golgi acidification, much less is known about cation efflux channels in the Golgi.

Human TMEM87A (hTMEM87A) and its homolog hTMEM87B, members of the Lung Seven Transmembrane Receptor (LUSTR) family^1^, are mainly present within Golgi. hTMEM87A is proposed to have a role in endosome-to-Trans Golgi Network (TGN) retrograde transport ^2,5^ which is also supported by its structural similarity with the Wnt transporter protein, Wntless^15^. However, a fraction of hTMEM87A localizes to the plasma membrane in cancer cells such as melanoma and glioblastoma multiforme (GBM), where it is associated with mechanosensitive cation channel activity^4,16^. Interestingly, in our accompanying study, we found that hTMEM87A which we renamed as GolpHCat, mediates voltage-gated, pH-sensitive, inwardly rectifying currents in both heterologous expression systems and reconstituted proteoliposomes and that various cation ions such as Cs^+^, K^+^, and Na^+^ can permeate through hTMEM87A. Furthermore, we found in TMEM87A KO mouse disorganized Golgi morphology, altered protein glycosylation/trafficking, and impaired hippocampal memory. Consequently, we concluded that hTMEM87A is critical for cellular functions and ion/pH homeostasis in the Golgi. However, its ion conduction and channel gating mechanism remained unclear. Here, we use single-particle cryo-electron microscopy (cryo-EM), molecular dynamic simulations, and whole-cell patch-clamp electrophysiology to gain structural and mechanistic insight into hTMEM87A as a voltage-gated non-selective cation channel.

## Overall structure of the human TMEM87A

Using Expi293F cells, we over-expressed full-length human TMEM87A (hTMEM87A, M1~E555), with the C-terminus fused to a thrombin cleavage sequence, EGFP, and a Twin-strep tag. After solubilization with n-dodecyl β-D-maltoside (DDM) and cholesteryl hemisuccinate (CHS), we purified hTMEM87A by Strep-Tactin affinity purification and size exclusion chromatography (SEC) in lauryl maltose neopentyl glycol (LMNG) and CHS (**Extended Data Fig.1a**). Since proteolytic cleavage of the C-terminal GFP and the Twin-strep tag with thrombin caused the partial precipitation of hTMEM87A protein, we used the homogeneous peak fractions containing hTMEM87A-EGFP-Twin-strep protein without proteolytic cleavage for structural studies and the analysis of *in vitro* channel activity. SDS-PAGE analysis of hTMEM87A after PNGase F treatment showed a shift to a lower molecular weight, indicating that hTMEM87A contains multiple N-linked oligosaccharides (**Extended Data Fig.1a**). Using single-particle electron cryo-microscopy (cryo-EM), we determined the structures of hTMEM87A at an overall resolution of 3.1 Å (**Fig.1a; Extended Data Fig.1, and Extended Data Table 1**). hTMEM87A contains GYG motif which is a signature selectivity filter of tetrameric K^+^ channels such as KcsA and HCN^17,18^. Unexpectedly, our SEC purification data and cryo-EM structure found that hTMEM87A is a monomer, not a tetramer. In the final density maps, we reliably assigned most of the side chains and all seven transmembrane segments as well as three N-linked oligosaccharides (N62, N79, and N127) (**Extended Data Fig.3**). Due to these glycans, we defined the globular domain containing N62, N79, and N127 as extracellular/luminal domain (ELD). We were not able to resolve two loops (L148~K167 and S193~L202) and the C-terminal tail (L474-E555 for both cases) in the cryo-EM map, probably due to their structural flexibility (**Fig.1b; Extended Data Fig.3**).

**Fig.1.**
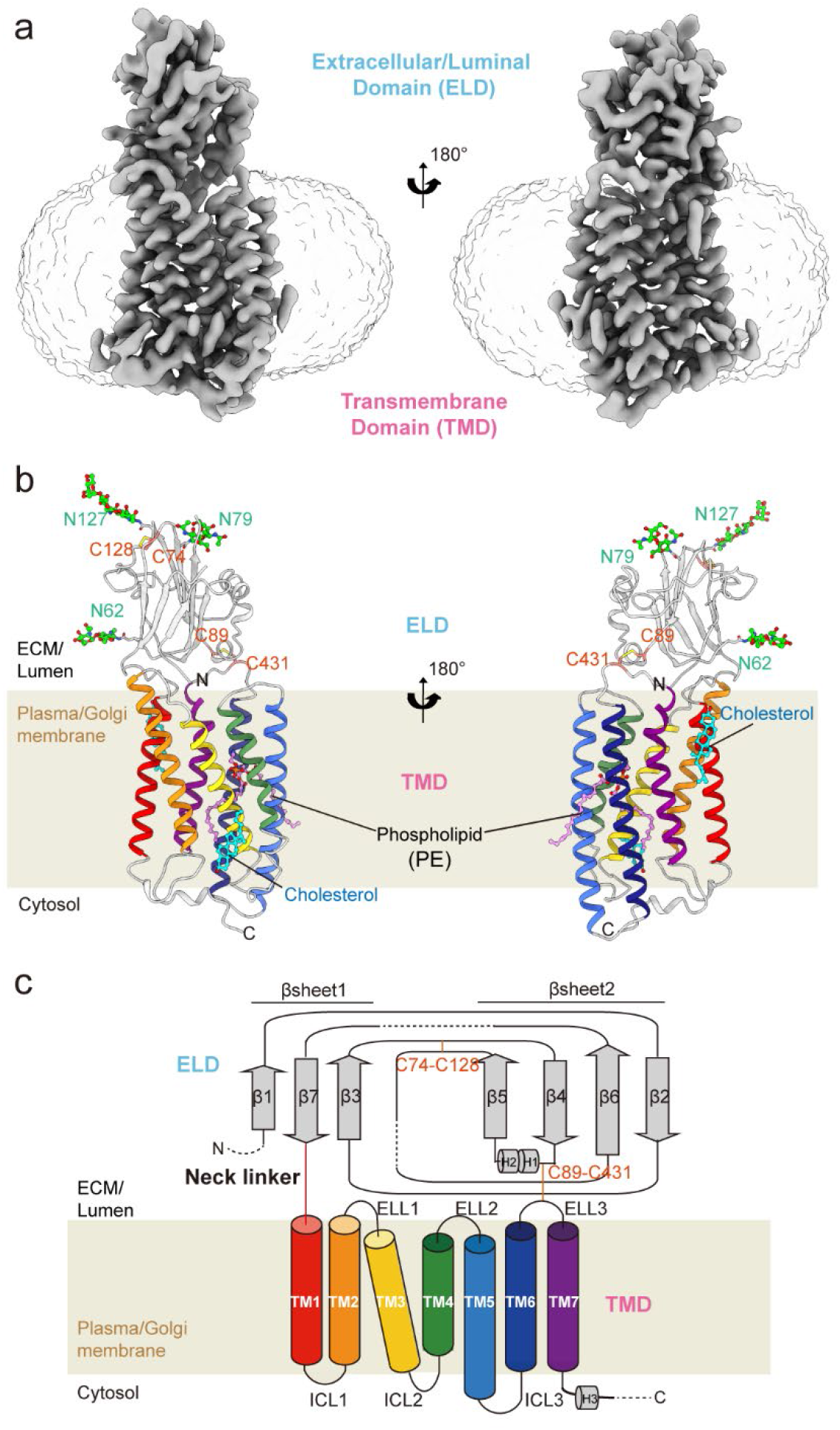
Cryo-EM Structure of human TMEM87A. **a,** Cryo-EM density map of the hTMEM87A. Extracellular/Luminal domain (ELD) and transmembrane domain (TMD) are colored in slate gray. The density of the detergent micelle (contoured at 0.161σ) is presented as light grey. **b,** Overall structure of hTMEM87A, with ELD colored in light grey and seven TMD helices in rainbow color from TM1 (red) to TM7 (purple). Disulfide bridges (orange, C74-C128, and C89-C431) and N-linked glycans (green, N62, N79, and N127) are shown as sticks. PE and cholesterol are indicated as pink and cyan sticks, respectively. **c,**Topology of hTMEM87A. ELD consists of two α-helices and seven β-strands arranged in an anti-parallel β-sandwich. The secondary structure elements (cylinder for helix and arrow for strand) are colored as in (b). Disulfide bonds are shown as an orange line. Dashed lines denote regions where density was insufficient for model building.

The seven transmembrane helices of hTMEM87A are arranged counterclockwise and connected by three intracellular loops (ICL1-ICL3) and three extracellular/luminal loops (ELL1-ELL3) (**Fig.1b-c**). Upstream of the transmembrane domains (TMD, Y224-P473), hTMEM87A has a globular domain (D38-K213) that spans either the extracellular or luminal compartments. This extracellular/luminal domain, hereafter referred to as ELD, resembles a β-sandwich fold connected to the first TMD helix by a linker (termed a neck linker, G214-D223). Interestingly, in the TMD pocket, we observed a well-resolved electron density that seemed to correspond to a phospholipid molecule, even though no phospholipids had been added during protein preparation (**Fig.1b; Extended Data Fig.3d**). Based on the shape of the electron density and the lipid compositions of the Golgi apparatus membrane^19^, we modeled phosphatidylethanolamine (PE; 1-palmitoyl-2-oleoyl-sn-glycerol-3-phosphoethanolamine, or PE-16:0-18:1) to this density. Two additional lipid-like densities around the TMD were modeled as a cholesterol, based on the shape of density. Taken together, these results indicate that hTMEM87A has a monomer architecture composed of ELD and seven TMD with a well-resolved PE in the TMD pocket.

## Structural features of the hTMEM87A ELD and PE-bound TMD

The hTMEM87A ELD is composed of seven anti-parallel β-strands (β1, β3, β7 for βsheet1 and β2, β4, β5, β6 for βsheet2) and two short α-helices (H1 and H2) (**Fig. 2a and Extended Data Fig.4a**). The helix-turn-helix motif (H1 and H2) located between β4 and β5 of ELD interacts with the outer surface of βsheet2 through long-range hydrophobic networks (I93, F96, V101, Y104, L105 and L108 on H1 and H2; Y52, Y83, Y118, I183 and I185 on βsheet2) and electrostatic interaction (K85-E92) (**Fig. 2a**). In addition, two conserved cysteines (C74 and C128) form a disulfide bond which bridges a long loop between β5 and β6 to a neighboring loop between β3 and β4 (**Fig. 2a**). In the TMD, TM1, TM6, and TM7 form a flat plane perpendicular to the lipid membrane (hereafter referred to as TM plane). The ELD makes broad interactions with the top edges of the flat TM plane and the extracellular loop connecting TM2 and TM3 (ELL1) (**Fig.2b and 2c, interaction patch 1~4**). Combined with the inter-domain disulfide bond (C89-C431), these broad interactions are likely to restrain ELD’s movement, thus maintaining a fixed orientation of ELD relative to the TM plane.

**Fig.2.**
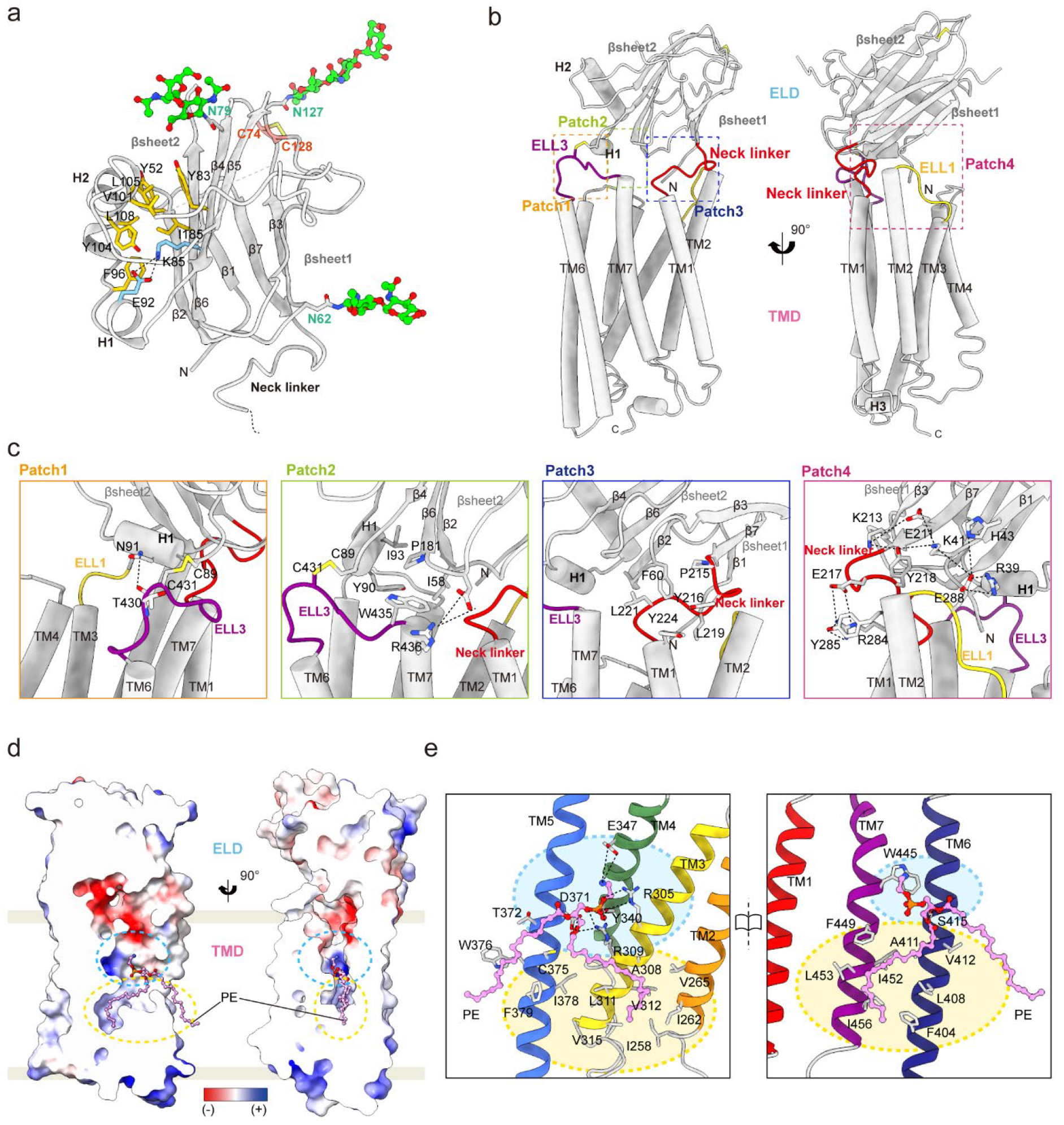
Structural features of the human TMEM87A ELD and PE-bound TMD. **a,** Structure of hTMEM87A ELD. Detail interactions of two short α-helices (H1 and H2) with the outer surface of βsheet2. Key interacting residues are displayed as sticks and labeled (ionic interaction: sky blue, hydrophobic interactions: yellow). N-linked glycans (N62, N79, and N127) are shown as green sticks. **b,** Interaction interfaces between ELD and TMD. Four interaction patches are indicated by dotted box (patch1, orange; patch2, lime; patch3, blue; and patch4, magenta). Loops involving interface interactions were highlighted (Neck linker: red, ELL1: yellow, and ELL3: purple). **c,** Close-up views of each interaction interface in **Fig. 2b**. Key interacting residues are shown as sticks and labeled. Hydrogen and ionic bonds are indicated as a dashed line. **d,** Two different views of the vertical cross-section of the PE-binding pocket in hTMEM87A TMD. View (left) is the same as in **Fig.1b, left**. PE is shown as a pink stick. The electrostatic surface potential of the central cavity is shown. The upper hydrophilic and lower hydrophobic cavities are indicated as cyan and yellow dashed circles, respectively. **e,** Open-book views of the PE-binding pocket and the interaction details. Interaction residues with PE are shown as sticks. The hydrogen and ionic bond are depicted as a dashed line. Cyan and yellow colored circle represent hydrophilic and hydrophobic cavities in hTMEM87A TMD, respectively.

We were intrigued to find a central cavity structure buried deep in the TMD. This cavity sits between the TM plane and the tilted/twisted TM2-TM5 helices and form two cavities; an upper hydrophilic cavity (cyan dashed circle) and a lower hydrophobic cavity (yellow dashed circle) (**Fig. 2d**). The upper hydrophilic cavity opens to the luminal side near the ELD and is exposed to the upper leaflet of the lipid bilayer through a gap between TM5 and TM6. However, its lateral side is partly sealed by PE, of which the R2-fatty acid chain also fills the lower hydrophobic cavity (**Fig. 2d**). Particularly, the terminal amine of PE forms a charge interaction with TM4 E347 and TM7 W445, and its phosphate group is coordinated by R305 and R309 of TM3, Y340 of TM4 and D371 of TM5 (**Fig. 2e**). In addition, oxygens in the ester group are stabilized by TM5 D371 and TM6 S415. Side chains of TM helices [α2 (I258,I262, and V265), α3 (A308, L311, V312, and V315), α5 (I378), α6(F404, L408, A411, and V412) and α7 (F449, I452, L453, and I456)] make hydrophobic interactions with the R2-fatty acid chain. The remaining R1-fatty acid chain of PE protrudes through the gap between TM5 and TM6, where it interacts with C375, W376, and F379 of TM5 and also likely with other lipid molecules in the membrane bilayer. The residues partaking in these extensive interactions with PE are highly conserved (**Fig. 2e and Extended Data Fig.5)**, indicative of a physiological role for PE in the structural and functional integrity of hTMEM87A.

## Comparison of hTMEM87A with its structural homologs

To investigate the structural basis of the TMEM87A ELD and TMD for its physiological roles we initially searched for structural homology of the hTMEM87A ELD using the Dali server^20^. We found that the β-sandwich domain of the hTMEM87A ELD resembles a ‘Golgi Dynamics’ (GOLD) domain present in p24 family proteins and SEC14-like protein 3 (PDB: 5AZW^21^, 5GU5^22^ and 4UYB, **Fig.3a and Extended Data Fig.6a**), which are implicated in the secretory pathway such as cargo sorting and membrane trafficking^23,24^. Interestingly, the sequence identity of the ELD across hTMEM87 family members is relatively low compared to that of the TMD (Sequence identity for ELD and TMD between hTMEM87A and hTMEM87B: 24.7 % and 63.1 %, respectively). However, the result from structure prediction with AlphaFold^25^ indicates that the β-sandwich domain and helix-turn-helix motif in ELD are highly similar between hTMEM87A and hTMEM87B (**Extended Data Fig.4b-c**), suggesting functional redundancy between TMEM87 family members. Given the roles of GOLD domain-containing proteins in the secretory pathway, the ELD of hTMEM87 family members may have a role in protein trafficking by interacting with yet unidentified partners. A comparison of amino acid sequences of TMEM87A and eukaryote orthologs^26^ also suggests that the evolutionary conserved TMD is more relevant to the physiological function of TMEM87A (**Extended Data Fig.5**).

**Fig.3.**
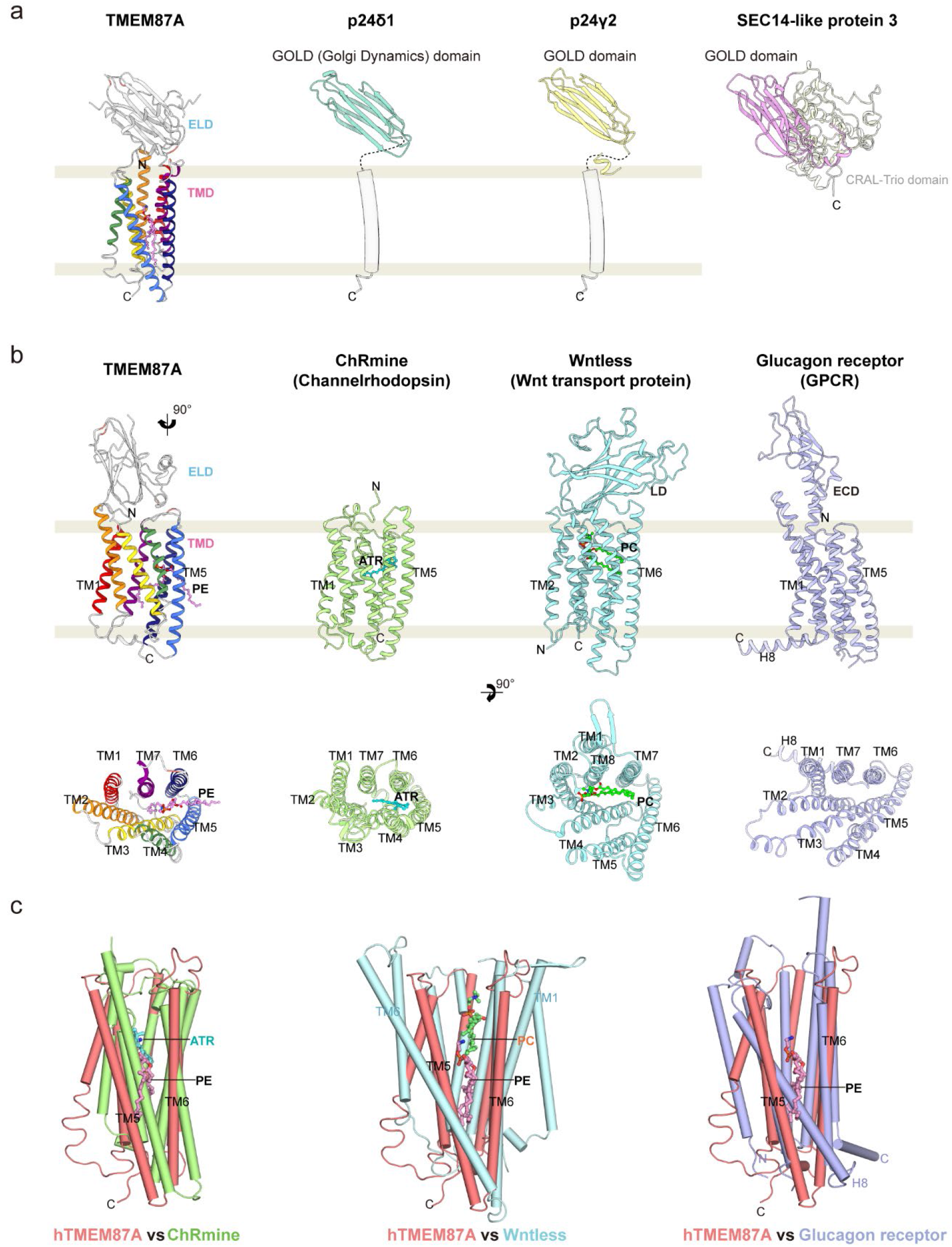
Comparison of hTMEM87A with its structural homologs. **a,** Structural comparison of the hTMEM87A ELD with other Golgi dynamics (GOLD) domain-containing proteins [p24δ1 (PDB: 5AZX, light teal), p24γ2 (PDB:5GU5, pale yellow), and SEC14-like protein 3 (PDB: 4UYB, light pink). **b,** Structural comparison of the hTMEM87A TMD with other seven transmembranes (7TM) proteins [channelrhodopsin (ChRmine, PDB: 7W9W, pale green), WNT transport protein (Wntless, PDB: 7DRT, cyan), and GPCR (Glucagon receptor, PDB: 5YQZ, light purple)], shown as a side view (top) and top view (bottom). The side view of hTMEM87A is the same as in **Fig.1b, left.** In the top view, ELD (D38-L221) of hTMEM87A, luminal domain (LD, P39-H225) of Wntless, extracellular domain (ECD, Q27-Q122) of glucagon receptor are omitted for clarity. Phosphatidylethanolamine (PE) in hTMEM87A, all-trans-retinal (ATR) in CHRmine, and Phosphatidylcholine (PC) in Wntless are displayed as pink, cyan, and lime sticks, respectively. **c,** Superimposition of hTMEM87A TMD and TM region of ChRmine (pale green), Wntless (cyan), or glucagon receptor (light purple), displayed as cartoons (cylinder for helix and arrow for strand). The view is a 90° rotation view of Fig. 3b along the y-axis to show the lateral opening between TM5 and TM6 of hTMEM87A TMD.

From a Dali server search^20^ with the hTMEM87A TMD, we found that structural homologs of the hTMEM87A TMD included the microbial channelrhodopsin (ChRmine protomer, PDB: 7W9W, TM1-TM7 among seven TM helices, Z-score 13.7, RMSD 3.3 over 214 residues), the Wnt transport protein (Wntless, PDB: 7DRT, TM2-TM8 among eight TM helices, Z-score 14.2, RMSD 4.1 over 233 residues), and the glucagon G-protein coupled receptor (Glucagon receptor, PDB: 5YQZ, TM1-TM7 among eight TM helices, Z-score 11.7, RMSD 4.0 over 223 residues)^27–29^ (**Fig.3b and Extended Data Fig.6b**), although overall sequence identity is relatively low (8~13%). hTMEM87A is similar to Wntless and the glucagon receptor in that a central cavity buried in the TMD is located below the globular extracellular domain and opened to the luminal side. However, the position of their extracellular domains differs substantially from that of the hTMEM87A ELD. Moreover, TM4 and TM5 of hTMEM87A are tilted toward the cavity core, resulting in a smaller central cavity compared to the cavities of Wntless and glucagon receptors that can accommodate a lipidated Wnt3a/8a hairpin and a peptide ligand, respectively (**Fig.3c**). The overall arrangement of hTMEM87A 7TM is much closer to that of ChRmine, although it does not have a globular extracellular domain. ChRmine and hTMEM87A are superimposed with an overall RMSD of 3.3Å over 7TM (214 residues) and two of their corresponding TMD layers (TM1/6/7 and TM2/3/4/5) are more tightly packed with each other than those of Wntless and the glucagon receptor (**Fig.3b**). Similar to all-*trans*-retinal in ChRmine, a PE molecule is surrounded by TM3-TM7 in hTMEM87A, even though their binding orientations are quite different (**Fig.3b and c**). Taken together, seven transmembrane helices of hTMEM87A are structurally similar to channelrhodopsin ChRmine, even though its extracellular globular domain resembles Golgi Dynamics domain.

## Putative Ion conduction pathway of hTMEM87A

Since voltage-gated inwardly rectifying currents were detected in proteoliposomes reconstituted with hTMEM87A, we examined the configuration of the hTMEM87A cavity using 3V^30^, a cavity, channel and cleft volume calculator, to delineate the location and shape of the ion-conducting pathway. A continuous channel pore was not observed in our structure (**Fig.4a**). In particular, the water-accessible cavity was physically blocked by the R2-fatty acid chain of PE, which extends from the lateral opening between TM5 and TM6 to the lower hydrophobic cavity. These data suggest that the observed conformation is presumably a closed state. However, analysis of electrostatic surface potentials for hTMEM87A revealed that negatively charged residues (D38 in ELD and E222, D223, E279, E298, D441, and D442 in TMD), main-chain carbonyl groups of L438 and W439, and the hydroxyl group of ELD Y90 are distributed on the funnel-shaped luminal vestibule (**Fig.4a-c,** hereafter negatively-charged luminal vestibule (NLV)). Like the extracellular vestibule of ChRmine^29^, the electronegative surface potential of the hTMEM87A NLV may attract and stabilize positively charged ions and thereby effectively increase the local cation concentration. To examine the potential role of these negatively charged residues of NLV in ion conduction, we performed mutagenesis and measured the hTMEM87A channel activity using whole-cell patch clamping. Among the three negatively charged residues, E279A mutation resulted in almost complete elimination of channel activity, whereas E298A mutation showed partially decreased channel activity and D442A mutation showed no change compared to WT (**Fig.4d, and Extended Data Fig.7a and 7b**). To further investigate the role of NLV in cation attraction, we performed accelerated Gaussian molecular dynamics (GaMD) simulations, which is an enhanced sampling technique widely used to explore biomolecular processes^31,32^. The GaMD simulations (3 replicas of 500 ns each) revealed that potassium cations (K^+^) are recognized by the superficial region of NLV (D38, Y90, E222, and D223), stabilized in deeper NLV regions upon interaction with additional residues (D441 and D442), and able to progress a short distance through the channel (**Fig.4e and Extended Data Fig.8a**). In fact, we found that K^+^ binding events occur frequently, as we explored up to 12 bindings within the 1.5 μs GaMD simulations. Collectively, cations attraction for ion conduction are orchestrated by the negatively charged-residues on the funnel-shaped luminal vestibule, while we did not observe complete cation permeabilization, possibly because we started with a closed structure and the expected timescale of the channel pore opening is long (~10 milliseconds).

**Fig.4.**
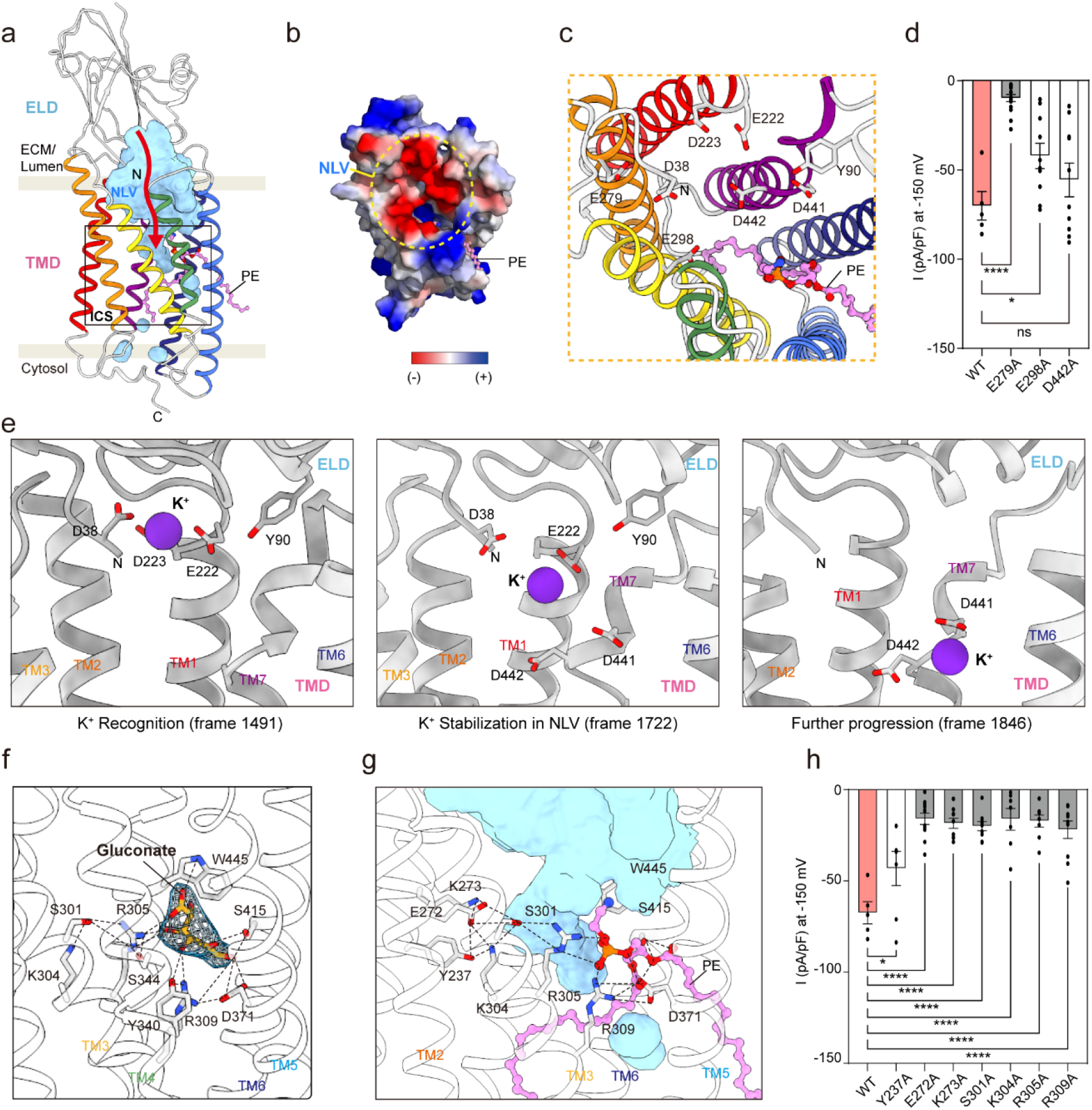
Putative ion-conducting pathway in hTMEM87A. **a,** Organization of ion-pathway in hTMEM87A. Water-accessible cavities are shown as a cyan surface in the hTMEM87A structure, with the putative ion conduction pathway indicated by a red arrow. Negative charged luminal vestibule (NLV) and constriction site (CS, black-lined box) are labeled. **b,** The surface electrostatic potential of the luminal vestibule. NLV is indicated as a yellow dotted circle. The hTMEM87A ELD and ten residues (R351-S361) of ELL2 are omitted for clarity. **c,** Close-up views of NLV and key negative-charged residues are shown as sticks. **d,** Current density measured at −150 mV for hTMEM87A WT and NLV mutants. Data are represented as mean ± SEM (n=5 for WT, n=15 for E279A, n=10 for E298A, and n=10 for D442A); one-way ANOVA with Dunnett’s test. ns> 0.005, *P<0.1, ****P<0.0001. **e**, Representative structures of the K+ conformational dynamics in the NLV of hTMEM87A obtained from GaMD simulations 1. K+ atoms are depicted as purple spheres, and their interacting residues as light grey sticks. TM4 and TM5 are omitted for clarity. **f**, Close-up view of gluconate binding site in hTMEM87A. Gluconate is displayed as a yellow stick with cryo-EM density map (contour level=0.118). Interaction residues are shown as grey sticks. The hydrogen and ionic bond are depicted as a dashed line. Helices of hTMEM87A TMD are displayed as transparent cartoons. **g**, Close-up views of CS and the interaction details. Key interaction residues and PE are shown as grey and pick sticks, respectively. The hydrogen and ionic bond are depicted as a dashed line. Cavities are shown as cyan surfaces. Helices of hTMEM87A TMD are displayed as transparent cartoons. TM4 is omitted for clarity. **h**, Current density measured at −150 mV for hTMEM87A WT and CS mutants. Data are represented as mean ± SEM (n=5 for WT, n=7 for Y237A, n=11 for E272A, n=9 for K273A, n=9 for S301A, n=7 for K304A, n=8 for R305A, and n=10 for R309A); one-way ANOVA with Dunnett’s test. *P<0.1, ****P<0.0001.

We have reported in accompanying paper that gluconate can effectively block both outward and inward currents of hTMEM87A with 0.1μM IC_50_. Thus, to decipher the binding pocket for gluconate and the putative Ion conduction pathway of hTMEM87A, we incubated purified hTMEM87A (0.7 mg/ml) with 10mM of sodium gluconate for 1hr on ice prior to freezing grids, and determined the cryo-EM structure of hTMEM87A-Gluconate (hTMEM87A-Gluc) at ~3.6 Å resolution (**Extended Data Fig.2**). While the overall structure of hTMEM87A-Gluc was essentially identical to hTMEM87A structure, of potential significance, the observed density indicates that a gluconate ion occupies the hydrophilic cavity of hTMEM87A via electrostatic interactions with R305, R309, D371 and W445, in a manner similar to the head group of PE (**Fig.4f and 4g, and Extended Data Fig.2h**). From these observations, we hypothesized that the extended electrostatic-interaction networks (Y237, E272, K273, S301, K304, R305, R309, S344, D371, Y340 and S415) underneath the NLV, which also mediate the binding of PE head group, can form a constriction and hence be implicated in hTMEM87A-mediated ion conduction upon the channel-opening stimulus. To test this hypothesis, we expressed hTMEM87A mutants (Y237A, E272A, K273A, S301A, K304A, R305A, and R309A) in CHO-K1 cells and recorded their respective channel activities by whole-cell patch clamp. Despite their robust cell membrane expression, current amplitudes at −150mV were significantly decreased for all mutants compared to wild type (**Fig. 4h, and Extended Data Fig.7d and 7e**). Moreover, these residues in the constriction sites are highly conserved (**Extended Data Fig.5**). Taken together, these data suggest that a channel-opening stimulus (high voltage in our experiments) can trigger rearrangements in electrostatic-interaction networks underneath the NLV, that ultimately lead to ion conduction. Conceivably, such a mechanism could resemble that of channelrhodopsin’s photocurrents, that are initiated by retinal isomerization which drive conformational changes of transmembrane helices^33–35^.

## Role of phosphatidylethanolamine and TM3 in ion conduction of hTMEM87A

As phosphatidylcholine (PC) is the most abundant lipid in the Golgi membrane (PC, ~50%; PE, ~20%)^19^, we next asked whether PC can compete with PE for binding to hTMEM87A. To this end, we compared the binding free energy of each lipid to hTMEM87A using MD simulations and assessed in a linear interaction energy (LIE) model^36^ [five 1 μs trajectories were simulated in the solvated system (S_p-L_), where hTMEM87A structures containing either lipid (PC or PE) bound to the TMD cavity were embedded in a simple Golgi model membrane (PC:PE=3:1)] (**Extended Data Fig.8b-8f**). The results showed that the van der Waals interaction energy of PC is increased due to the additional methyl groups, while its coulombic interaction energy decreased because of their screening effect. However, the calculated binding free energy of PC to hTMEM87A is higher than that of PE by 19.2 kJ/mol (ΔF_m->p_ (PC) = −13.2 kJ/mol and ΔF_m->p_ (PE) = −32.4 kJ/mol), suggesting that under physiological condition, PE is more likely to bind to hTMEM87A. Importantly, these data support that the PE observed in the cryo-EM structure is co-purified with hTMEM87A from the Golgi membrane.

Next, we sought to characterize the entry of PE and binding to hTMEM87A. To this end, we simulated five trajectories up to 1 μs starting from a system (named Sp*/L) constructed by removing the bound PE lipid (L) from the hTMEM87A cryo-EM structure and then embedding the bare hTMEM87A (p*) in a lipid bilayer composed of only PE. In order to characterize the binding process, we define two distances, dP and dR2-Cent, as those from the phosphorus atom and the center of mass of PE’s R2-fatty acid chain to the smallest-moment principal axis (Pz) of TMD, respectively (**Fig. 5a**). In 2D histogram of dP and dR2-Cent for all five trajectories, we identified seven highly populated states of PE (from S1 to S7), among which the fully bound state (S1) is most probable. The conformational snapshots that correspond to these seven states are shown in **Fig. 5b**. In one example trajectory, for which variations of dP and dR2-Cent as a function of time are presented, PE lipid starts entering the TMD cavity quickly (~25 ns) and goes through two intermediate partially bound states (S3 and S4) to arrive at the fully bound state (S1) after ~500 ns (**Fig. 5c**). These data demonstrate how lipid entry can occur as a stepwise process. Moreover, the closeness between the state corresponding to the cryo-EM structure (S*) and S1 indicates a clear tendency for PE to become inserted at that unique position of hTMEM87A. Of note, at the end of the five simulation trajectories, PE either becomes fully bound (two cases) or becomes stuck in metastable unbound states (S5-7, three cases). It has been previously reported that changes in membrane tension can cause displacement of lipid molecules in mechanosensitive ion channels such as MscS and TRAAK, leading to conformational changes and channel opening^37–39^. Therefore, we asked whether the PE displacement in hTMEM87A could be responsible for a channel opening, in a similar manner as other mechanosensitive ion channels. To this end, we designed a S415F mutant that should block PE entry to the central cavity of the hTMEM87A TMD through the lateral gap between TM5 and TM6, and then measured channel activity. S415F mutant displayed significantly larger current amplitudes at −150 mV compared to WT (**Fig. 5d**), suggesting that the inhibition of PE insertion into hTMEM87A TMD increases open probability. Taken together, these observations indicate that, while the fully bound state observed in the cryo-EM structure is unique and stable, there are several partially bound and unbound states of PE, which is critical for the opening of ion-conducting pathway.

**Fig.5.**
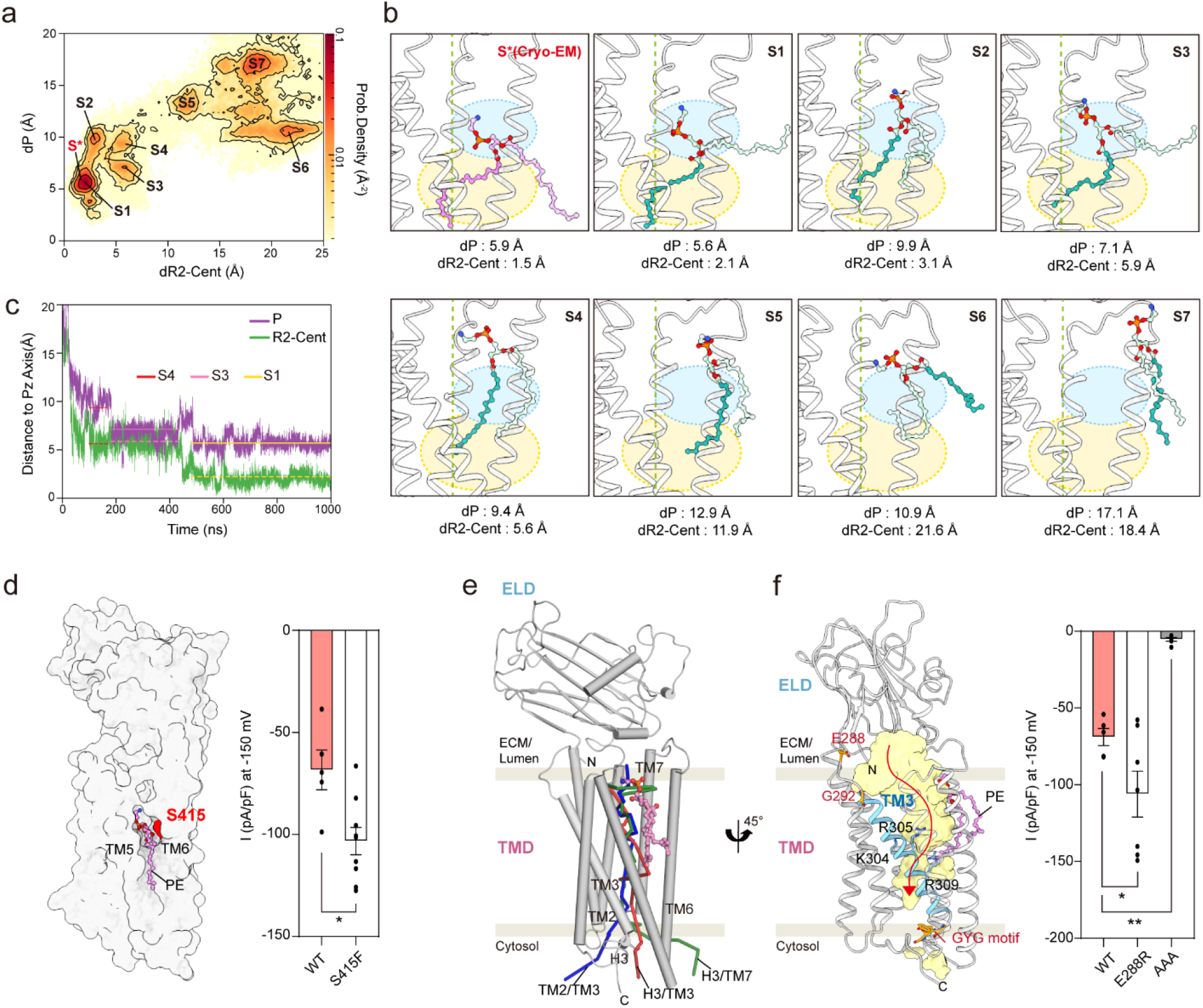
MD simulation of PE entry and role of PE in hTMEM87A channel activity. **a,** 2D histogram of distances (dP and dR2-Cent) for five MD trajectories (5×1 μs from system S_p*/L_). dP and dR2-Cent are the distances from PE’s phosphorus atom and the center of mass of PE’s R2-fatty acid chain to the smallest-moment principal axis (Pz) of TMD, respectively. PE position in our cryo-EM structure is indicated as red, and seven highly populated states of PE are labeled (from S1 to S7). **b,** The conformational snapshots of PE in seven different states (S1-S7). PE from cryo-EM structure (pink) and MD simulations are displayed as sticks. The phosphorus atom, R1- and R2-fatty acid chain of PE from MD simulations are colored as orange, light blue, and teal, respectively. Dashed green lines indicate the smallest-moment principal axis (Pz) of TMD. TM4 and TM5 (P324-R392) are omitted for clarity. Calculated distances of P and R2-Cent are indicated. **c,** Variations of distances of dP and dR2-Cent as a function of time along one trajectory. States of S4, S3, and S1 are indicated by horizontal lines. **d,** Current density measured at −150 mV for hTMEM87A WT and lateral-opening block mutants. Data are represented as mean ± SEM (n= 5 for WT and n=9 for S415F); t-test. *P<0.05. **e,** Representative water permeation pathway. A structure of hTMEM87A in S5 from Fig.5b was used for calculation and displayed as cartoons (cylinder for helix and arrow for strand). Representative of three groups for possible water permeation pathway [H3/TM3 (4 cases, path5 as a representative), TM2/TM3 (6 cases, path14 representative), H3/TM7 (10 cases, path23 representative)] are indicated as red, blue and green, respectively. **f,** Organization of possible ion-pathway in hTMEM87A. Water-accessible cavities are shown as a yellow surface in the hTMEM87A structure (S5 from Fig.5b), with the putative ion conduction pathway indicated by a red arrow. Potential voltage-sensing motif TM3 and three positive charged residues (K304, R305, and R309) are colored as cyan. Residues corresponding to the mutants (E288R and located at either end of TM3 (E288R on ELL1 and GYG (318-320) on ICL2 ->AAA) are indicated by orange sticks. The current density was measured at −150 mV for hTMEM87A WT and mutants (E288R and AAA). Data are represented as mean ± SEM (n= 5 for WT, n=7 for E288R, and n=6 for AAA); one-way ANOVA with Dunnett’s test. *P<0.1, **P<0.01.

In the accompanying study, we show that the channel activity of hTMEM87A is regulated in a voltage-dependent manner. Voltage-regulated ion channels, such as K_v_ channels, Na_v_ channels, and trimeric intracellular cation (TRIC) channels, usually have a voltage-sensing domain (helix) wherein positively charged lysine or arginine residues are enriched^40–43^. Particularly, the TRIC-B1 channel contains three conserved basic residues on its TM4 helix. These interact with the phosphate group of phosphatidylinositol-4,5-biphosphate (PIP2) and nearby negatively charged residues, which occlude its ion permeation pathway. Moreover, the artificial disulfide bonds that locks the voltage-sensing TM4 helix of TRIC in a restrained conformation causes it to remain closed upon depolarization, thus demonstrating an essential mechanism in which the voltage-sensing TRIC TM4 helix is coupled to channel activation^41^. Although the structure of hTMEM87A is different from TRIC channels, three conserved basic residues (K304, R305, and R309) lining one side of the hTMEM87A TM3 interacts with an electronegative PE head group and the neighboring E272 residue, which closely resemble the voltage-sensing TM4 helices of TRICs (**Fig. 5f**). Reportedly, the different mechanosensitive channel activity between human and mouse TMEM87A are attributed to two non-conserved amino acids (L271 and G292 for human; F271 and N292 for mouse), among which the flexible G292 residue is located at the beginning of the TM3 helix^4^. Moreover, we found here that mutations in loop residues flanking the TM3 helix (E288R on ELL1 or GYG [318-320] on ICL2 ->AAA) affected hTMEM87A ion conduction (**Fig. 5f**). In addition, from MD simulation using one trajectory of S_p*/L_ that sampled the S5 PE-unbound state (see **Fig. 5a**) most of the time (**Fig. 5e**), we identified that there are three possible water pathways [H3/TM3(4 cases), TM2/TM3 (6 cases), H3/TM7 (10 cases), 8 outliers from total 28 calculated water pathways] and that water molecules exit through a small hole close to the cytoplasmic tip of TM3 helix. Collectively, these results suggest a critical role for the hTMEM87A TM3 in influencing channel activity, probably through participating in voltage-sensing and gating. However, we did not observe any conventional channel structure from the geometric pore analysis and cation permeation pathway among all simulated trajectories (~1 μs). Possible explanations for the lack of cation permeation pathway may include 1) longer simulation times (>1 μs) are needed, and 2) the lack of structures for open state upon the voltage stimulation. Future work is needed to test these possibilities.

The TMEM87 family is widespread in the eukaryotic organism and predominantly localized in the Golgi membrane. In this work, we determined the high-resolution cryo-EM structures of hTMEM87A in complex with the Golgi membrane phospholipid PE and the pharmacological inhibitor gluconate. Because the domain arrangement of hTMEM87A closely resembles that of Wntless, a regulator of Wnt protein sorting and secretion, and other GOLD domain seven-transmembrane helix (GOST) proteins^15^, the most parsimonious conclusion is that hTMEM87A is involved in the trafficking, secretion, and sorting of yet unidentified proteins. While future studies will be required to examine its function, our structure analysis of hTMEM87A, complemented by MD simulations and electrophysiological analysis, provide crucial insights into mechanistic aspects of hTMEM87A as a non-selective voltage-gated cation channel. Although residues on the funnel-shaped luminal vestibule can in principle attract cations, the R2-fatty acid chain of PE and key residues that interact with its head group appear to occlude ion conduction pathway, indicating that hTMEM87A under cryo-EM condition (resting state) is impermeable to ions. Notably, the putative ion conduction pathway is lined with largely hydrophobic and electropositive residues that are highly conserved. Although currently unclear, we speculate that conformational changes on the voltage-sensor TM3 of hTMEM87A and PE displacement may occur cooperatively during gating in response to membrane depolarization under physiological conditions, similar to the gating mechanism of TRICs, that eventually leads to the opening of the ion conduction pathway of hTMEM87A. Further structural and functional studies are required to address the gating dynamics of the TMEM87 family and their broader physiological roles.

## Methods

### Cloning, protein expression and purification

cDNA encoding human TMEM87A (hTMEM87A, NP_056312.2, M1-E555) followed by a TEV protease cleavage sequence (ENLYFQG), a PreScission Protease cleavage sequence (LEVLFQGP), EGFP (M1-239K), a thrombin cleavage sequence (LVPRGS) and a Twin-strep-tag were cloned into the BamHI and XhoI sites of a pcDNA3.4 (#A14697, Invitrogen). All hTMEM87A mutants were created by site-directed mutagenesis using the WT construct as a template. Constructs and primer information are listed in **Extended Data Tables 2 and 3.** For the whole-cell patch clamp, the coding sequence of hTMEM87A WT or mutants (M1-E555) was cloned into the SalI and BamHI sites of pIRES2-Dsred vectors (#632463, addgene).

Recombinant hTMEM87A proteins (WT and mutants) were transiently expressed in Expi293F cells (#A14527, Thermo Fisher Scientific) according to the manufacturer’s instructions. Briefly, 200 μg of plasmid DNA was transfected into 200 ml of Expi293F cells (3.0 × 10^6^ cells/ml) using Expifectamine (#A14524, Thermo Fisher Scientific). Cells were cultured in Expi293 expression medium (#A14351, Thermo Fisher Scientific) at 37°C and 8% CO2 with shaking (orbital shaker, 120 rpm). After 20 h, the enhancer (#A14524, Thermo Fisher Scientific) was supplemented to the culture, then further incubated for 30-34 hours at 30°C. Cell pellets were resuspended in 20 ml TN buffer [50mM Tris pH 9.0, 250mM NaCl, and 1x complete protease inhibitor cocktail (#11836170001, Roche)] and lysed by sonication (total 2min, 1sec with intervals of 5sec, 20% amplitude). After ultracentrifugation (Beckman Ti70 rotor, 150,000 × g for 1 h), the collected membrane fraction was homogenized with a glass Dounce homogenizer in 20ml buffer [TN buffer + 1% (w/v) *n*-dodecyl β-D-maltoside (DDM; #D310S, Anatrace) and 0.2% (w/v) cholesteryl hemisuccinate (CHS; #CH210, Anatrace)] and solubilized for 2h at 4°C. The insoluble cell debris was removed by ultracentrifugation (Beckman Ti70 rotor, 150,000 x g for 1 h), and the supernatant was incubated with 2ml Strep-Tactin resin (#2−1201-025, IBA Lifesciences) for 30 min at 4°C. After washing with 10 column volumes of wash buffer [TN buffer + 0.05% (w/v) lauryl maltose neopentyl glycol (LMNG; #NG310, Anatrace) and 0.01% (w/v) CHS], the hTMEM87A-EGFP-Twin strep tag was eluted with 5ml elution buffer [TN buffer + 10mM desthiobiotin]. After concentration using Amicon Ultra centrifugal filter (100-kDa cut-off; Millipore), the hTMEM87A-EGFP-Twin strep was further purified by size exclusion chromatography (SEC) using a Superose 6 Increase 10/300 GL column (#29-0915-96, Cytiva) equilibrated with a final buffer [TN buffer + 0.01% (w/v) LMNG and 0.002% (w/v) CHS]. The collected peak fractions were concentrated to ~0.8 mg/ml using Amicon Ultra centrifugal filter (100-kDa cut-off; Millipore) and immediately used for the cryo-EM grid preparation for the hTMEM87A structure.

For the complex cryo-EM structure of hTMEM87A with gluconate (hTMEM87A-Gluc), HEPES buffer was used instead of Tris buffer during purification (HN buffer; 50mM HEPES pH 7.5, 250mM NaCl, and 1x complete protease inhibitor cocktail). Other protein solubilization and purification conditions were the same as those for hTMEM87A. Before freezing grids, 10mM of sodium gluconate (S2054, Sigma) was added to purified hTMEM87A (0.7 mg/ml) and incubated for 1hr on ice.

### Cryo-EM sample preparation and data collection

Quantifoil R 1.2/1.3 Cu 200-mesh holey carbon grids (#4220C-XA, SPI SUPPLIES) were glow-discharged for 75 s at 15 mA (PELCO easiGlow Glow Discharge Cleaning system, Ted Pella). Then, 4 μl of the purified hTMEM87A or hTMEM87A-Gluc were applied to the grid at 100% humidity at 4°C. After 7 s blotting, grids were plunged into liquid ethane using a FEI Vitrobot Mark IV (ThermoFisher Scientific). Micrographs were acquired on a Titan Krios G4 TEM operated at 300 keV with a K3 direct electron detector (Gatan) at the Institute for Basic Science (IBS), using a lit width of 20 eV on a GIF-quantum energy filter. EPU software was used for automated data collection at a calibrated magnification of ×105,000 under the single-electron counting mode and correlated-double sampling (CDS) mode^44^, yielding a pixel size of 0.849 Å/pixel. The micrograph was dose-fractionated to 57 frames under a dose rate of 7.95 e^−^/pixel/sec with a total exposure time of 6.14 s, resulting in a total dose of about 67.72 e^−^/ Å^2^. A total of 10,377 movies for hTMEM87A and 13,099 movies for hTMEM87A-Gluc were collected with a nominal defocus range from −0.8 to −1.9 μm. Detailed parameters were summarized in **Extended Data Table 1**.

### Cryo-EM data processing, model building and refinement

The detailed image processing workflow and statistics are summarized in **Extended Data Fig. 1c and 2c and Extended Data Table 1**. Micrographs were subjected to patch motion correction and patch CTF estimation in cryoSPARC v.3.3.2^45^. For the hTMEM87A data set, 96,330 particles were first picked using a blob picker of cryoSPARC. Then, 2D class average images were generated as templates for subsequent reference-based auto-picking. A total of 8,101,104 particles from the complete datasets were binned four times, and to identify higher quality particles, subsequent 2D classification, *Ab initio*, and heterogeneous refinement were performed in cryoSPARC. The resulting 445,198 particles from the 3D classes showing good secondary structural features were re-extracted into the original pixel size for further 3D refinements. Non-uniform refinement^46^ and CTF refinement^47^ improved the particle alignment and map quality. The final refinement yielded a map at an overall ~3.1 Å resolution according to the 0.143 cut-off criterion^48^. For hTMEM87A-Gluc, reference-based picked 6,035,205 particles were processed similarly to hTMEM87A data processing. The final non-uniform refinement from 201,915 particles yielded a map at an overall ~3.6 Å resolution. The mask-corrected Fourier shell correlation (FSC) curves were calculated in cryoSPARC, and reported resolutions were based on the gold-standard Fourier shell correlation (FSC) = 0.143 criteria. Local resolutions of density maps were estimated by Blocres^49^. Model building for hTMEM87A was initiated using the module ‘Map to model’ in PHENIX package^50^ and a model generated by AlphaFold^25,51^. The model was then subjected to iterative manual and automated refinement rounds in PHENIX and Coot^52^. The final refinement statistics are summarized in Extended Data Table 1. The final refinement statistics are summarized in **Extended Data Table 1**.

### Model analysis

A cavity search using Solvent Extractor from the Voss Volume Voxelator server^30^ was performed using an outer-probe radius of 5 Å and an inner-probe radius of 1.2 Å. The Dali server^20^ was used to search protein structures having a similar fold. All molecular graphics figures were prepared with UCSF ChimeraX^53^ and PyMOL^54^.

### Whole-cell patch clamp in CHO-K1 cells overexpressing hTMEM87A WT or mutants

Although endogenous human TMEM87A mainly localized to the Golgi membrane, overexpressed hTMEM87A is partially localized to the plasma membrane^4^ (**Extended Data Fig.7e).**Therefore, whole-cell patch clamping was performed to measure the channel activity of hTMEM87A localized at the plasma membrane. 2.5 μg of hTMEM87A WT or mutant-pIRES2-DsRed was transiently transfected into CHO-K1 cells (2×10^5^ cells/well in a six-well plate) on the day before patch recording. After 24 hours, cells were seeded onto 0.1 mg/ml PDL-coated coverslips in 24 wells and used for whole-cell patch clamp recording within 12 hours. The bath solution contained 150mM NaCl, 3mM KCl, 10 mM HEPES, 5.5mM glucose, 2mM MgCl_2_, 2mM CaCl_2_ with pH adjusted to pH7.3 by NaOH (320~325 mOsmol/kg). Borosilicate glass pipettes were pulled with a micropipette puller (P-97, Sutter Instrument) and had a resistance of 4–6 MΩ in the bath solution when filled with a pipette solution containing 130mM K-gluconate, 10mM KCl, 10mM HEPES, 10mM 1,2-Bis(2-amino phenoxy)ethane-*N*,*N*,*N*’,*N*’-tetraacetic acid (BAPTA) with pH adjusted to pH7.3 by KOH (290~310mOsmol/kg). The holding voltage was −60mV. For recording with voltage-clamp ramp protocol, currents were measured under the 1000 ms-duration voltage ramps descending from +100 mV to −150 mV with 10 s time intervals. Electrical signals were amplified using MultiClamp 700B (Molecular Devices, USA). Data were acquired by Digitizer 1550B (Molecular Devices, USA) and pClamp 11 software (Molecular Devices, USA) and filtered at 2 kHz.

### Surface biotinylation assay and western blot

The biotinylation assay was performed using the Pierce^™^ Cell Surface Protein Biotinylation and Isolation Kit (Thermo Scientific, A44390) as manufacturer’s instructions. In detail, WT hTMEM87A or its point mutants (E279A, E298A, D442A, Y237A, E272A, K273A, S301A, K304A, R305A, R309A, S415F, and S415W) cloned into the pIRES2-DsRed vector were transiently transfected into HEK293A cells (5×10^6^ cells in 100 mm dish). After 40 hours, cells were rinsed with PBS and then incubated with 0.25 mg/ml Ez-Link^™^-Sulfo-NHS-SS-Biotin in PBS for 10 min at room temperature. Biotinylated cells were washed with ice-cold TBS three times, and harvested cell pellets resuspended in 250 μl lysis buffer containing a complete protease inhibitor cocktail (Roche). After centrifugation (15,000 × g for 5 min at 4°C), the supernatant was collected, and protein concentration was measured using the Bradford protein assay. 800 μg of protein was incubated with 250 μl of NeutrAvidin agarose resin (Thermo Scientific, 29200) on a shaker for 30 min at room temperature. After three times washing using the washing buffer in the Pierce^™^ Cell Surface Protein Biotinylation and Isolation Kit, biotinylated proteins were eluted using 100 μl elution buffer containing 10mM DTT, separated by 10 % SDS–PAGE and transferred to PVDF membranes. The membranes were blocked with 5% skim milk in TBST for 1 hour at RT, and washed with TBST three times. Then, the membranes were incubated with primary antibodies: rabbit anti-TMEM87A antibodies (1:1000, NBP1-90531, Novus Biologicals) and mouse anti-β-actin antibodies (1:2000, sc-69879, Santa Cruz) overnight at 4°C. The membranes were washed with TBST three times and incubated with the corresponding horseradish peroxidase-conjugated secondary antibodies [HRP-linked anti-rabbit IgG (#7704, Cell Signalling) and HRP-linked anti-mouse IgG, #62-6520, Thermo Scientific] for 1 hour at RT. After washing three times with TBST, immune-reactive protein bands were detected using EzWestLumiOne (#2332632, ATTO).

### Gaussian Accelerated Molecular Dynamics (GaMD) simulations

Gaussian accelerated molecular dynamics (GaMD) is an unconstrained enhanced sampling method that smooths the potential energy surface and reduces the energy barriers of biomolecular processes by adding a harmonic boost potential^31,32^.

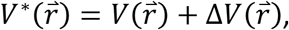

Where 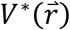 is the modified potential, 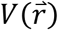 is the system potential and 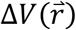 is the harmonic boost potential. Along the simulation time, the harmonic boost potential is only added when system potential drops below reference energy:

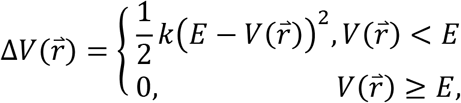

Where *E* is the reference energy, and *k* is the harmonic force constant. The two adjustable parameters *E* and *k* can be determined by applying the following criteria:

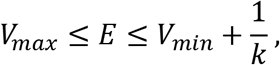

Where *V_max_* and *V_min_* are the maximum and minimum potential energies, respectively.

The PE-bound hTMEM87A structure was used as a starting point and minimized in a two-stage geometry optimization approach using Gaussian accelerated molecular dynamics (GaMD) simulation. First, a short minimization of the water molecules positions, with positional restraints on the protein, ligand, and P31 atoms of the membrane, was performed with a force constant of 10 kcal/mol Å^-2^ at constant volume periodic boundary conditions. Second, an unrestrained minimization including all atoms in the simulation cell was carried out. The minimized system was gently heated in two phases. First, the temperature was increased from 0K to 100K in a 20 ps step. Harmonic restraints of 10 kcal/mol Å^-2^ were applied to the protein, ligand, and membrane. Second, the temperature was slowly increased from 100K to the production temperature (310.15K) in a 100 ps step. In the second phase, harmonic restraints of 10 kcal/mol Å^-2^ were applied to the protein, ligand, and P31 atoms of the membrane. The Langevin thermostat was used to control and equalize the temperature. The initial velocities were randomized in the heating step. In the heating and following steps, bonds involving hydrogen were constrained with the SHAKE algorithm, and the time step was set at 2 fs, allowing potential inhomogeneities to self-adjust. The equilibration step was performed in three stages. First, 5 ns of MD simulation under NVT ensemble and periodic boundary conditions were performed to relax the simulation temperature. Second, 5 ns of MD simulation under NPT ensemble at a simulation pressure of 1.0 bar was performed to relax the density of the system. The semi-isotropic pressure scaling using the Monte Carlo barostat was selected to control the simulation pressure. Third, additional 5 ns of MD simulation was performed to relax the system further. A cutoff value of 11 Å was applied to Lennard-Jones and electrostatic interactions.

After equilibration, an extra short 5 ns of MD simulation followed by 45 ns GaMD simulation (the boost potential is applied) was carried out in order to collect potential statics for calculating the acceleration parameters. Finally, a 500 ns of GaMD production run was performed. In total, we performed 3 independent simulations (i.e., 1.5 μs accumulated time). All GaMD simulations were performed using the AMBER2021 package^55^ and applying the ‘dual-boost’ potential, where one boost potential is applied to the dihedral energetic term and the other to the total potential energetic term of the force field. The reference energy was set to the upper bound, which provides a more aggressive boost. The upper limit of the boost potential standard deviation, σ0, was set to 6.0 kcal/mol.

### Molecular Dynamics simulations and analysis for PE binding to hTMEM87A

All MD simulation systems were built from the cryo-EM structure of hTMEM87A (PDB ID: 8HSI) using Membrane Builder in CHARMM-GUI^56^. The missing loops (L148-K167 and S193-L202) were reconstructed using Modeller^57^. Using the PropKa program, the protonation state for K273 was determined as LYN, H187, and H403 as HID, and other histidine residues as HIE. All systems were solvated in ~150 mM KCl solution and the periodic simulation boxes were about 10 × 10 × 14 nm^3^ large. The Amber FF14SB, Lipid17, and TIP3P force fields were used for protein, lipid, and water, respectively^58,59^. The standard CHARMM-GUI equilibration protocol was followed to equilibrate the systems. The Particle-mesh Ewald method^60^ was used for the electrostatic interaction, and a cut-off length of 0.9 nm for the van der Waals interaction. The production trajectories were integrated with a time step of 2 fs using OpenMM^61^. The temperature was kept at 310.15 K via the Langevin dynamics with a friction coefficient of 1 ps−1, and the pressure was retained at 1 bar via a Monte Carlo barostat with a coupling frequency of 5 ps−1. The trajectory analysis was performed with MDAnalysis^62^, and the first 100 ns was discarded when calculating interaction energies.

From simulations of the lipid binding process (m->p), m-L + p* -> m + p-L, where m stands for membrane, we estimate the timescale for the hTMEM87A-PE binding process to be ~100 ns. We then try to estimate the timescale of the unbinding process by calculating the binding free energy *ΔF_m→P_*(*L*) of the hTMEM87A-lipid binding process according to the linear interaction energy (LIE) model^36^. For that purpose, we simulated five 1 μs trajectories for the solvated system (S_p-L_) of the cryo-EM structure embedded in a simple Golgi model membrane (m), which has a PC-to-PE ratio of 3:1, and one 1 μs trajectory for the solvated system (S_m-L_) that consists of only the membrane. From the p-L trajectories, we computed average coulombic and van der Waals interaction energies corresponding to the L->p process of L(g) + p* -> p-L as

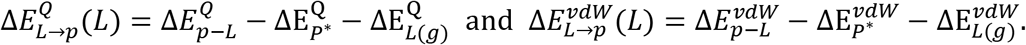

From the m-L trajectory, we similarly computed for the L->m process of L(g) + m -> m-L.

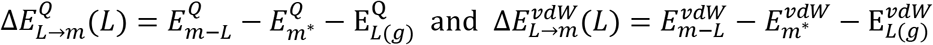

We can then obtain the average interaction energies for the m->p process by

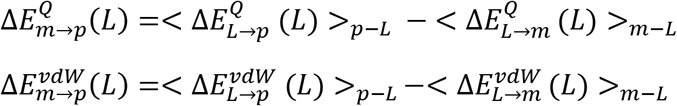

Finally, the binding free energy is estimated by the LIE formula of

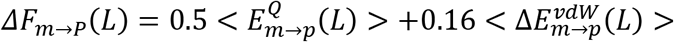

## Data Availability

The atomic coordinate for hTMEM87A and hTMEM87A with gluconate has been deposited to the Electron Microscopy Data Bank and the Protein Data Bank with the accession numbers PDB: 8HSI (EMD-34998) and PDB: 8HTT (EMD-35017), respectively

## Acknowledgments

We are grateful to the staff of the Research Solution Center at IBS for help with cryo-EM data collection. Computational work for this research was performed on the data analysis hub (Olaf) in the IBS Research Solution Center. This work was supported by grants from the Institute for Basic Science (IBS-R030-C1 to H.M.K. and IBS-R001-D2 to C.J.L). This work was also supported by the Mid-career Researcher Program (NRF-2020R1A2C2101636), Bio & Medical Technology Development Program (2022M3E5F3080873), and Brain Pool Program (NRF-2021H1D3A2A02038434, NRF-2021H1D3A2A02081370) funded by the Ministry of Science and ICT (MSIT) through the National Research Foundation of Korea (NRF). We also thank the Korea Institute of Science and Technology Information (KISTI) Supercomputing Center (KSC-2021-CRE-0469) and the Ewha Womans University Research Grant of 2022 (to S.C.).

## Author contributions

A-R.H. and H.M.K. designed the experiments and analyzed the data; A-R.H. purified proteins and determined the cryo-EM structure; A.Z., M.A.M.S., and S.C. performed MD simulation and analyzed the data; H.J.K. and C.J.L. performed electrophysiology experiments; and A-R.H., A.Z., M.A.M.S., H.J.K., C.J.L., S.C., and H.M.K. wrote the manuscript.

