## Extended Figure & Table for "Structural insights into ion conduction by novel cation channel, TMEM87A, in Golgi apparatus"

Extended Date Figures 1-8 and Legends

Extended Date Tables 1–3

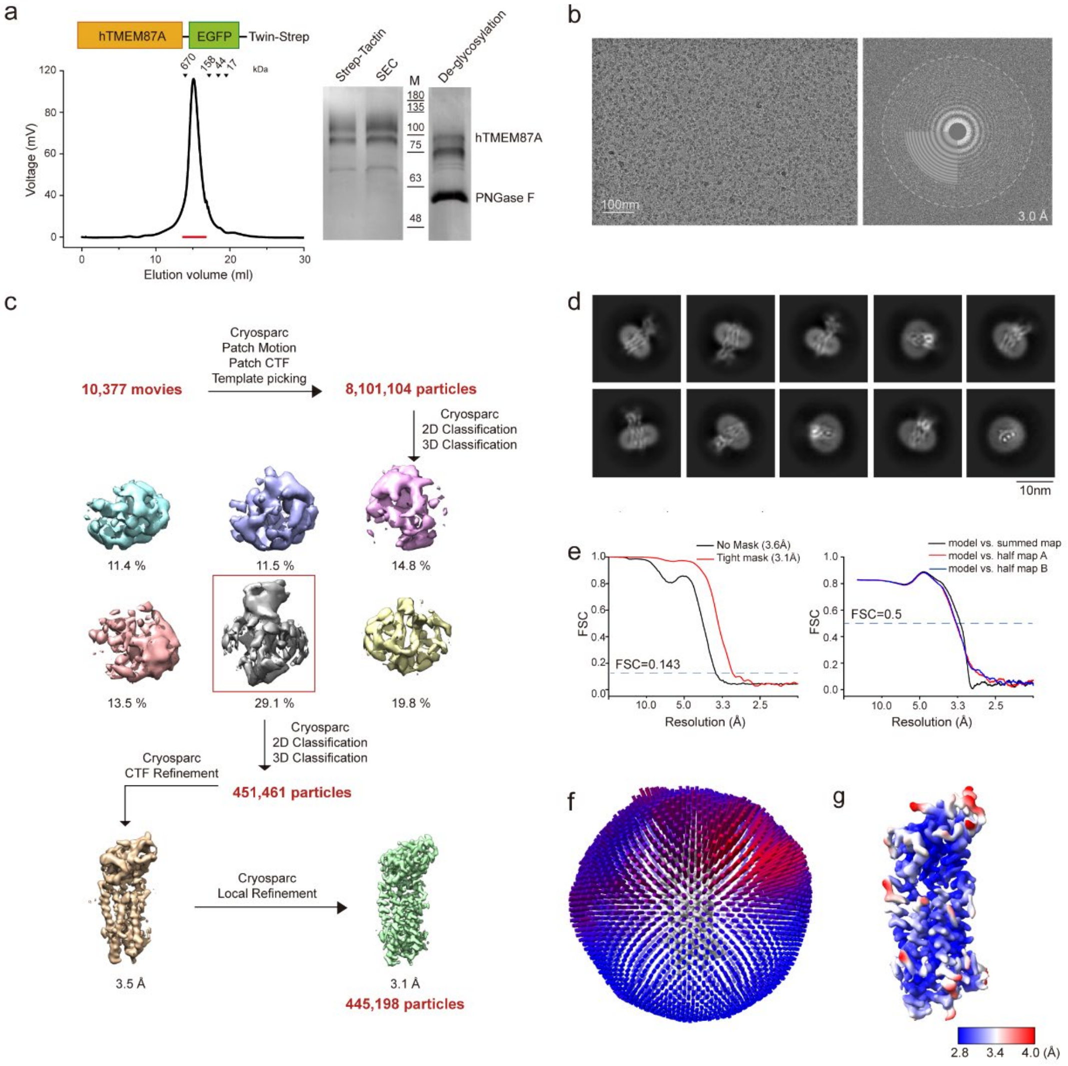


**Extended Data Fig.1 | Purification and Cryo-EM analysis of the hTMEM87A.**

**a**, Expression construct and size-exclusion chromatography (SEC) profile of hTMEM87A-EGFP-Twin-strep (left). Pooled fractions for cryo-EM analysis are marked with the red bar. Coomassie blue stained SDS-PAGE gel of the elution fractions from Strep-Tactin affinity purification and the SEC peak fractions, respectively (middle). SDS-PAGE analysis of hTMEM87A-EGFP-Twin-strep after PNGase F treatment (right).

**b**, Representative cryo-EM micrograph (left) and its Fourier transform (right).

**c**, Data processing workflow of cryo-EM analysis of hTMEM87A.

**d**, Representative 2D class averages of hTMEM87A.

**e**, Fourier shell correlation (FSC) curves between two independently refined half maps in cryoSPARC (left, resolution cutoff at FSC = 0.143) and FSC curves for cross-validation of a model (right, resolution cutoff at FSC = 0.5).

**f**, Euler angle distribution of all particles used in the final 3D reconstructions. The cylinder bars' height and color (from blue to red) are proportional to the number of particles in those views.

**g**, Final cryo-EM map of hTMEM87A-pH9 colored with local resolution.

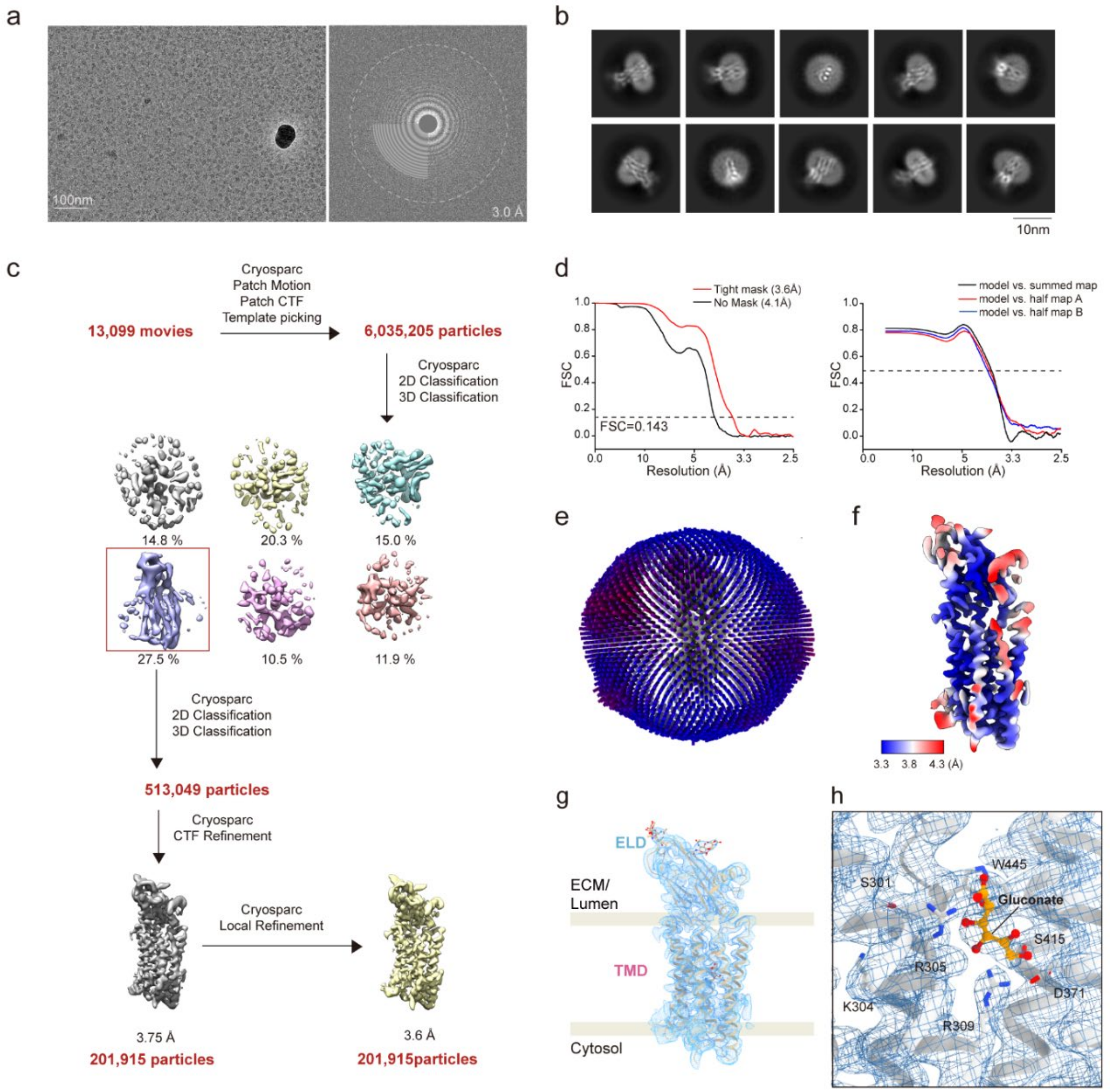


**Extended Data Fig.2 | Purification and Cryo-EM analysis of the hTMEM87A-Gluc.**

**a,** Representative cryo-EM micrograph (left) and its Fourier transform (right) of the hTMEM87A-Gluc.

**b,** Representative 2D class averages of hTMEM87A-Gluc.

**c,** Data processing workflow of cryo-EM analysis of hTMEM87A-Gluc.

**d,** Fourier shell correlation (FSC) curves between two independently refined half maps in cryoSPARC (left, resolution cutoff at FSC = 0.143) and FSC curves for cross-validation of a model (right, resolution cutoff at FSC = 0.5).

**e,** Euler angle distribution of all particles used in the final 3D reconstructions. The cylinder bars' height and color (from blue to red) are proportional to the number of particles in those views.

**f,** Final cryo-EM map of hTMEM87A-Gluc colored with local resolution.

**g,** Fit of hTMEM87A-Gluc model into 3.6Å cryo-EM map (contour level = 0.123).

**h,** Cryo-EM map and model of hTMEM87A-Gluc in Glutamate-binding site (contour level = 0.125). Glutamate and interacting residues are shown as sticks and labeled.

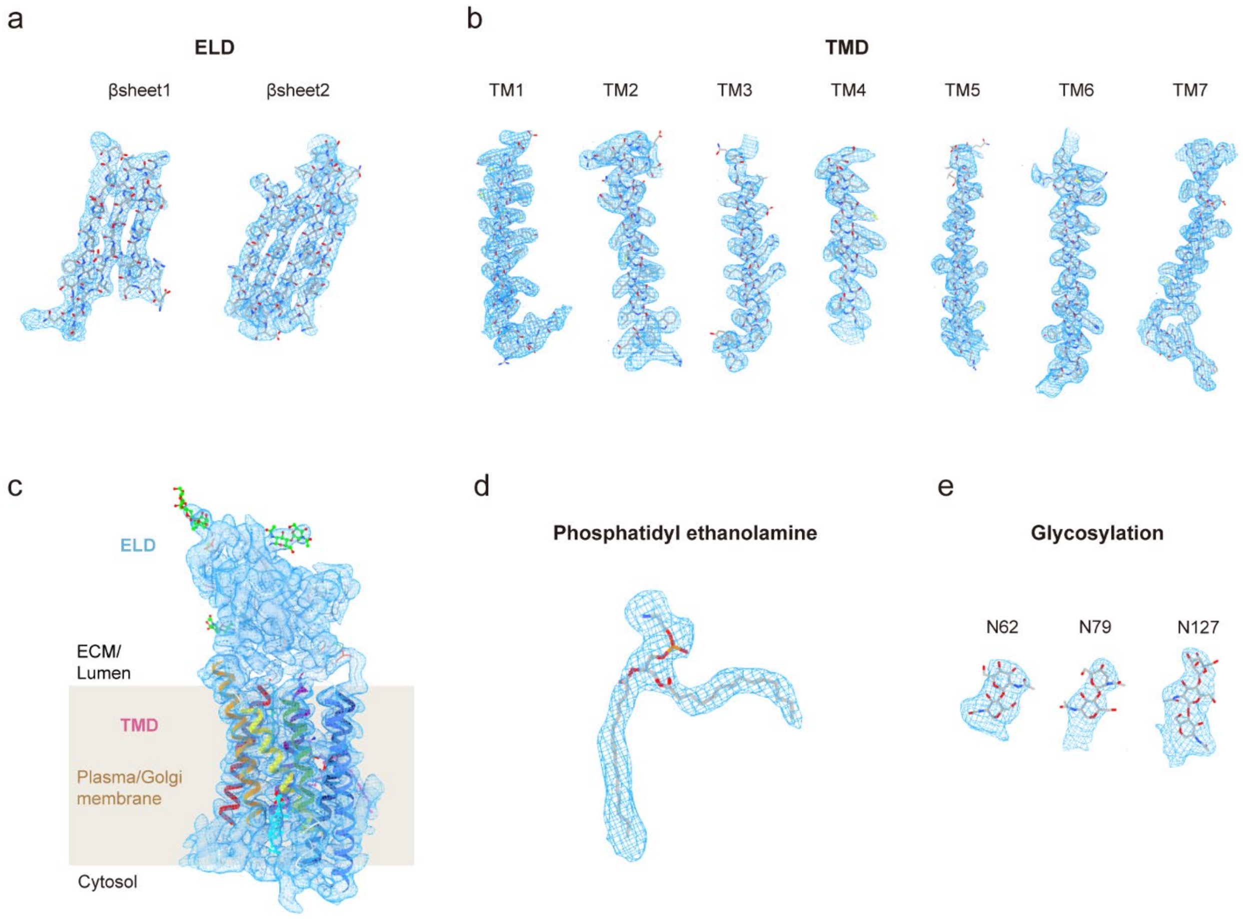


**Extended Data Fig.3 | Cryo-EM map for hTMEM87A.**

**a,** Cryo-EM map and model from hTMEM87A for βsheet1 (β1, D38-S48; β3, N62-E72; β7, N204-P215, contour level = 0.142) and βsheet2 (β2, G49-F60; β4, L76-D88; β5, L117-Q126, β6, A180-S192, contour level = 0.129) of ELD.

**b,** Cryo-EM map and model of seven helices of TMD (TM1, E222-L256, contour level = 0.101; TM2, R257-E288, contour level = 0.101; TM3, V290-V322, contour level = 0.101; TM4, V333-G354, contour level = 0.101; TM5, Q356-R393, contour level = 0.101; TM6, N394-I428, contour level = 0.101; and TM7, S433-P473, contour level = 0.101).

**c,** Fit of hTMEM87A model into 3.1Å cryo-EM map (contour level = 0.107). The hTMEM87A is displayed in cartoon and colored with the same scheme as in Fig.1b.

**d,** Cryo-EM map and model of the phosphatidylethanolamine (contour level = 0.101).

**e,** Cryo-EM map and model of three N-linked oligosaccharides (N62, contour level = 0.0805; N79, contour level = 0.0805; and N127, contour level = 0.0805).

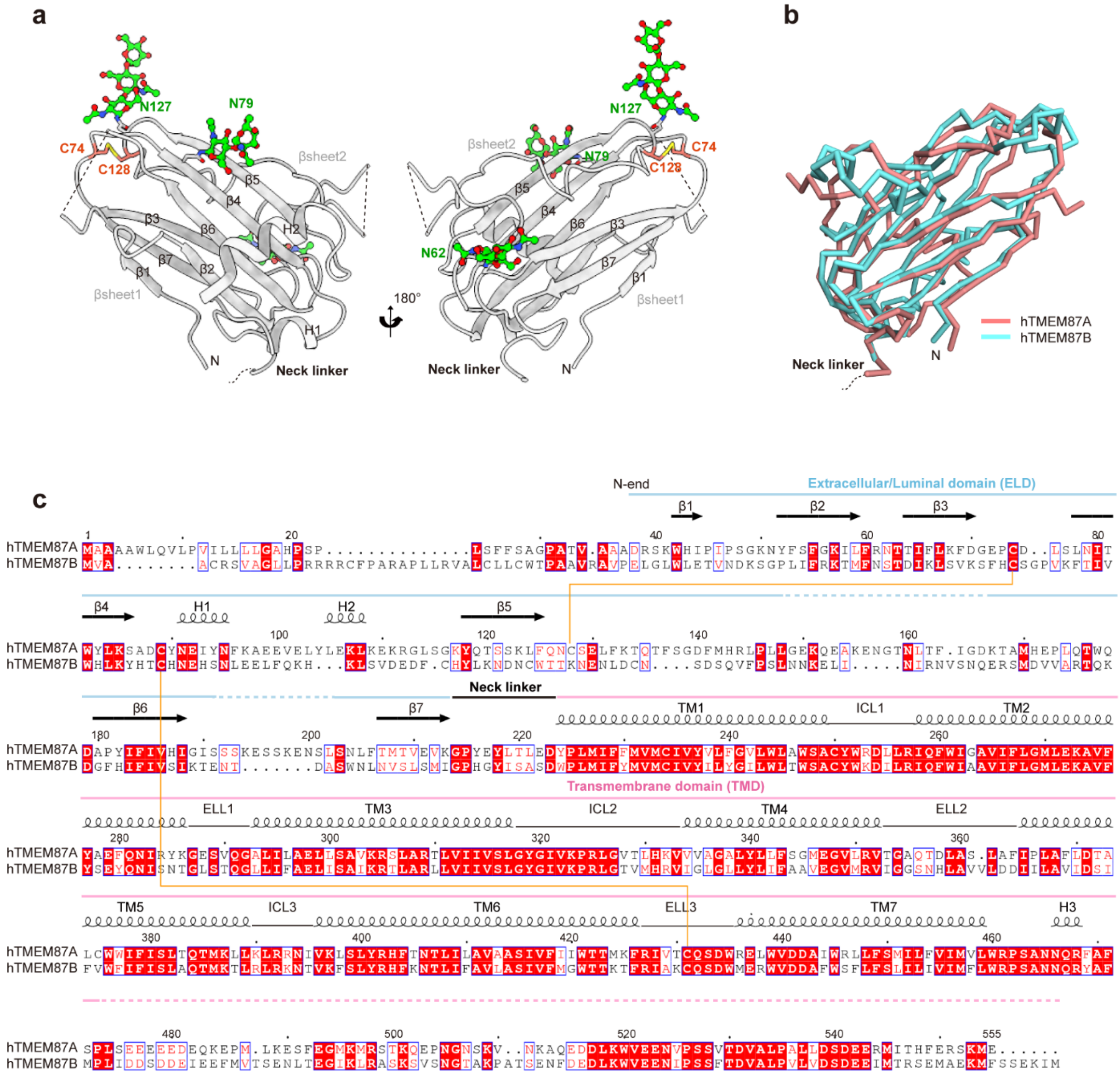


**Extended Data Fig.4 | Structure of hTMEM87A ELD and comparison with hTMEM87B ELD.**

**a,** Two different views of hTMEM87A ELD with anti-parallel β-sandwich fold (βsheet1 and βsheet2). Structures are shown as a cartoon and colored in light gray. The glycosylation site (N62, N79, and N127) and disulfide bridge (C74-128) are indicated. Disulfide bridge (yellow) and N-linked glycans (green) are shown as sticks. Disordered regions were indicated as dashed lines.

**b,** Superimposition of hTMEM87A ELD (salmon) with the hTMEM87B ELD (cyan, NP_116213.1, E45-S212), whose structure is predicted by Alphafold. The calculated Cα root mean square deviation (R.M.S.D) is 2.24.

**c,** Amino acid sequence alignment of hTMEM87A and hTMEM87B (NP_116213.1). Domains (ELD, TMD, N-end, and neck linker), secondary structure elements (arrows for β stands and helices for α-helices), and disulfide bond (orange lines) are displayed. Red boxes indicate perfect sequence conservation, whereas blue-lined boxes show residues with >70% similarity based on physicochemical properties. The sequence alignment was created using Clustal Omega and EsPript 3.0.

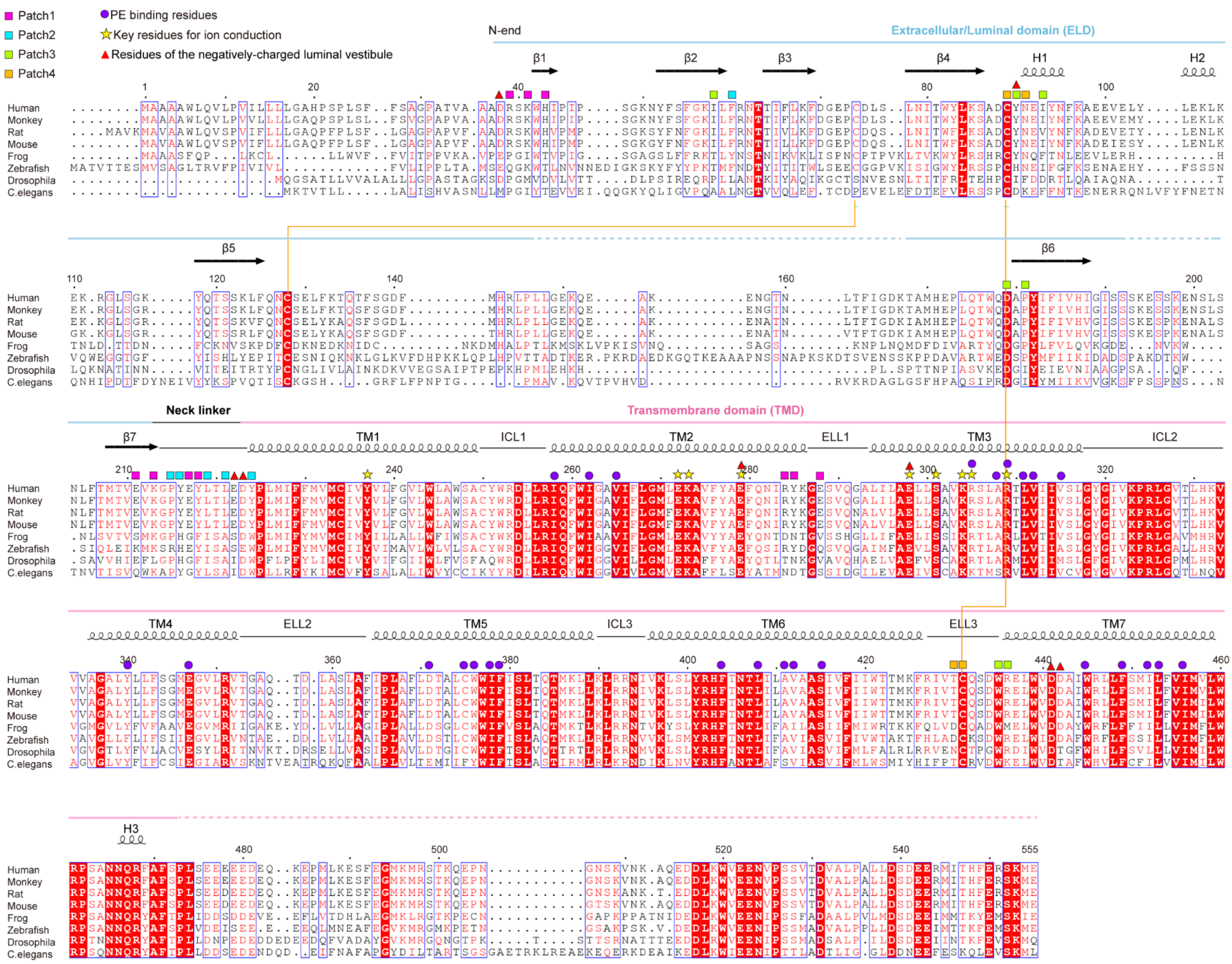

Patch1
Patch2
Patch3
Patch4
PE binding residues
Key residues for ion conduction
Residues of the negatively-charged luminal vestibule
N-end
Extracellular/Luminal domain (ELD)
β1
β2
β3
β4
H1
H2
β5
β6
β7
Neck linker
Transmembrane domain (TMD)
TM1
ICL1
TM2
ELL1
TM3
ICL2
TM4
ELL2
TM5
ICL3
TM6
ELL3
TM7
H3
Human
Monkey
Rat
Mouse
Frog
Zebrafish
Drosophila
C.elegans

**Extended Data Fig.5 | Multiple sequence alignment of TMEM87A**.

Amino acid sequence alignment of human (*Homo sapiens*, NP_056312.2), monkey (*Chlorocebus sabaeus,* XP_008015094.1), rat (*Rattus norvegicus*, XP_038962452.1), mouse (*Mus musculus*, NP_776095.2), frog (*Xenopus tropicalis*, NP_001016057.1), Zebrafish (*Danio rerio*, NP_001082853.2), Fruit fly (*Drosophila melanogaster*, NP_608612.3), Roundworm (*Caenorhabditis elegans*, NP_508729.2). TMEM87A orthologs are obtained by ortholog search[1]. Domains (ELD, TMD, N-end, and neck linker) and secondary structure elements (arrows for β stands and helices for α-helices) are displayed above the alignment. Squares indicate interacting residues at the interface between ELD and TMD (patch1: orange, patch2: green, patch3: cyan, and patch4: magenta). Purple circles indicate interacting residues with phosphatidyl ethanolamine (PE). Yellow stars indicate key residues involved in the ion conduction. Red triangles indicate residues consisting of the negatively-charged luminal vestibule. Disulfide bonds (orange lines) are also indicated. Red boxes indicate perfect sequence conservation, whereas blue-lined boxes show residues with >70% similarity based on physicochemical properties. The sequence alignment was created using Clustal Omega[2] and EsPript 3.0[3].

**a**

| Protein name | p24δ1 | p24γ2 | SEC14-like protein 3 |
|---|---|---|---|
| PDB ID | 5AZX | 5GU5 | 4UYB |
| Dali Z-score | 6.8 | 5.4 | 6.0 |
| RMSD | 2.5 Å over 85 residues | 2.5 Å over 78 residues | 3.1 Å over 93 residues |

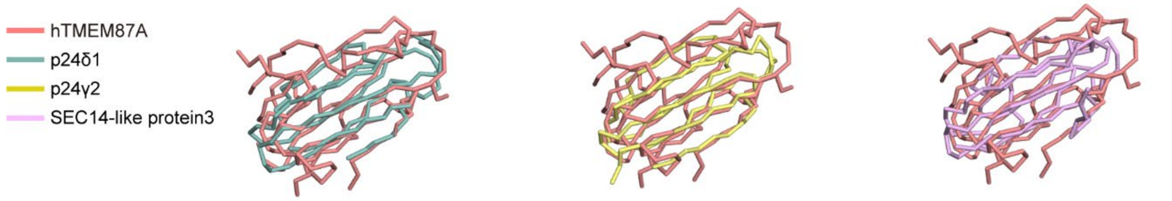


**b**

| Protein name | ChRmine | Wntless | Glucagon receptor |
|---|---|---|---|
| PDB ID | 7W9W | 7DRT | 5YQZ |
| Dali Z-score | 13.7 | 14.2 | 11.7 |
| RMSD | 3.3 Å over 214 residues | 4.1 Å over 233 residues | 4.0 Å over 223 residues |

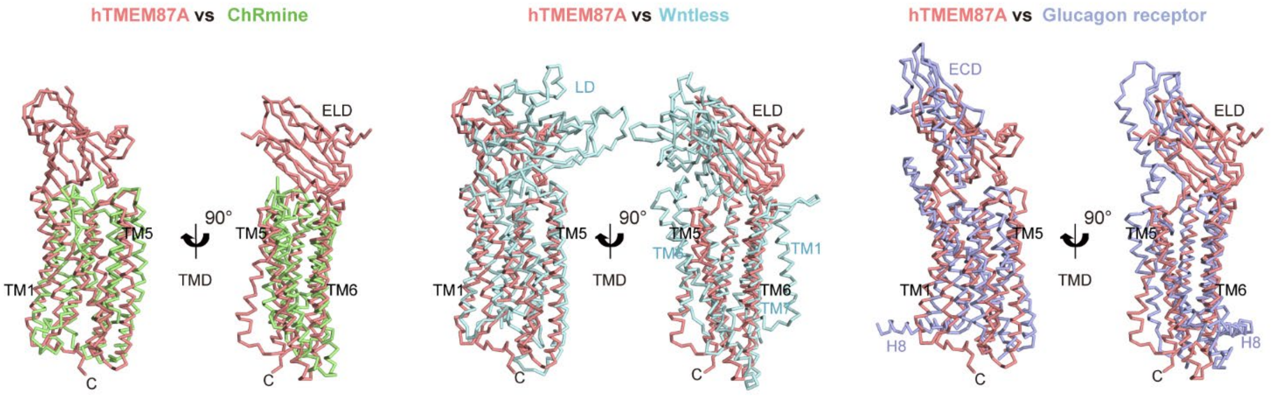


**Extended Data Fig.6 Structural homology of hTMEM87A.**

**a**, Structural homology search with Dali server, identifying that hTMEM87A ELD resembles the GOLD domain of p24δ1 (PDB: 5AZX), p24γ2 (PDB: 5GU5), and SEC14-like protein 3 (PDB: 4UYB). Result summary of Dali server search (Z-score and Cα RMSD) are presented. Superimposition of hTMEM87A ELD (salmon, A37-E222) and the GOLD domain of p24δ1 (G30-G129, light teal), p24γ2 (G31-K141, yellow), or SEC14-like protein 3 (K275-P386, light pink), shown as ribbon diagrams.

**b**, Structural homology search with Dali server, identifying that hTMEM87A TMD resembles seven transmembrane helices (7TM) of ChRmine (PDB:7W9W), Wntless (PDB:7DRT), and glucagon receptor (PDB:5YQZ). Result summary of Dali server search (Z-score and Cα RMSD) are presented. Superimposition of hTMTM87A TMD and TM region of ChRmine (TM1-7, pale green), Wntless (TM2-8, cyan), or glucagon receptor (TM1-7, light purple), shown as ribbon diagrams.

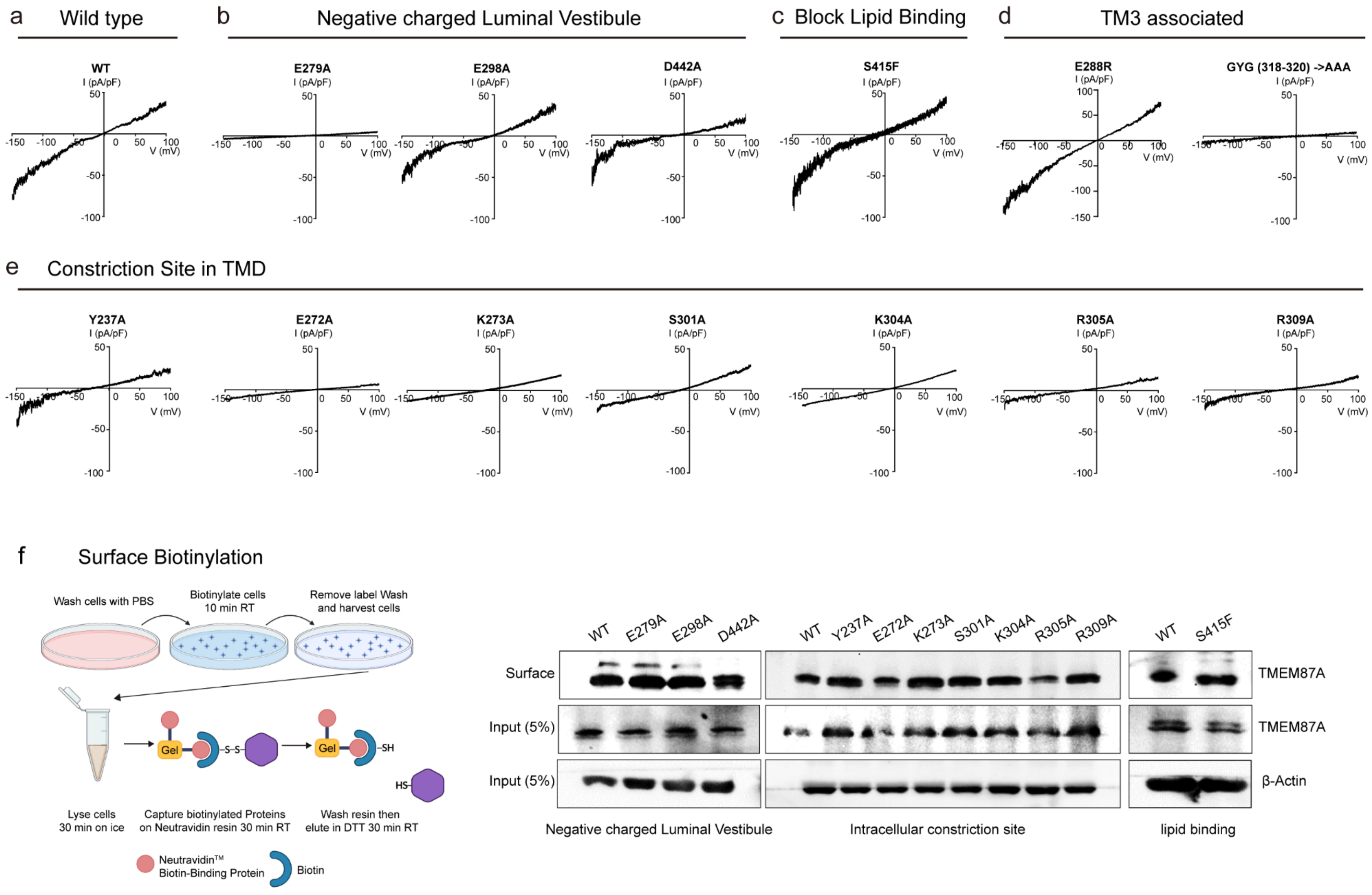


**Extended Data Fig.7 Electrophysiology of hTMEM87A mutants.**

**a-e**, Representative current-voltage (I-V) curves obtained in whole-cell configuration of hTMEM87A WT and its mutants. The currents were recorded with voltage-clamp ramp protocol descending from +100 mV to -150 mV.

**a**, Representative current-voltage (I-V) curves of hTMEM87A WT.

**b**, Representative current-voltage (I-V) curves of hTMEM87A mutants (E279A, E298A, and D442A) for NLV.

**c**, Representative current-voltage (I-V) curves of hTMEM87A mutants (S415F) for blocking lipid binding.

**d**, Representative current-voltage (I-V) curves of hTMEM87A mutants (E288R and AAA) for TM3 associated

**e**, Representative current-voltage (I-V) curves of hTMEM87A mutants (Y237A, E272A, K273A, S301A, K304A, R305A, and R309A) for constriction site in TMD

**f**, Membrane expression of hTMEM87A WT and its mutant measured by surface biotinylation assay. HEK293A cells were transfected with hTMEM87A WT or its mutants (E279A, E298A, D442A, Y237A, E272A, K273A, S301A, K304A, R305A, R309A, S415F, and S415W), treated with biotin to isolate surface hTMEM87A proteins. Proteins were immunoblotted with TMEM87A antibodies (Novus). These results indicate that all of hTMEM87A mutants are expressed in the plasma membrane at the similar level.

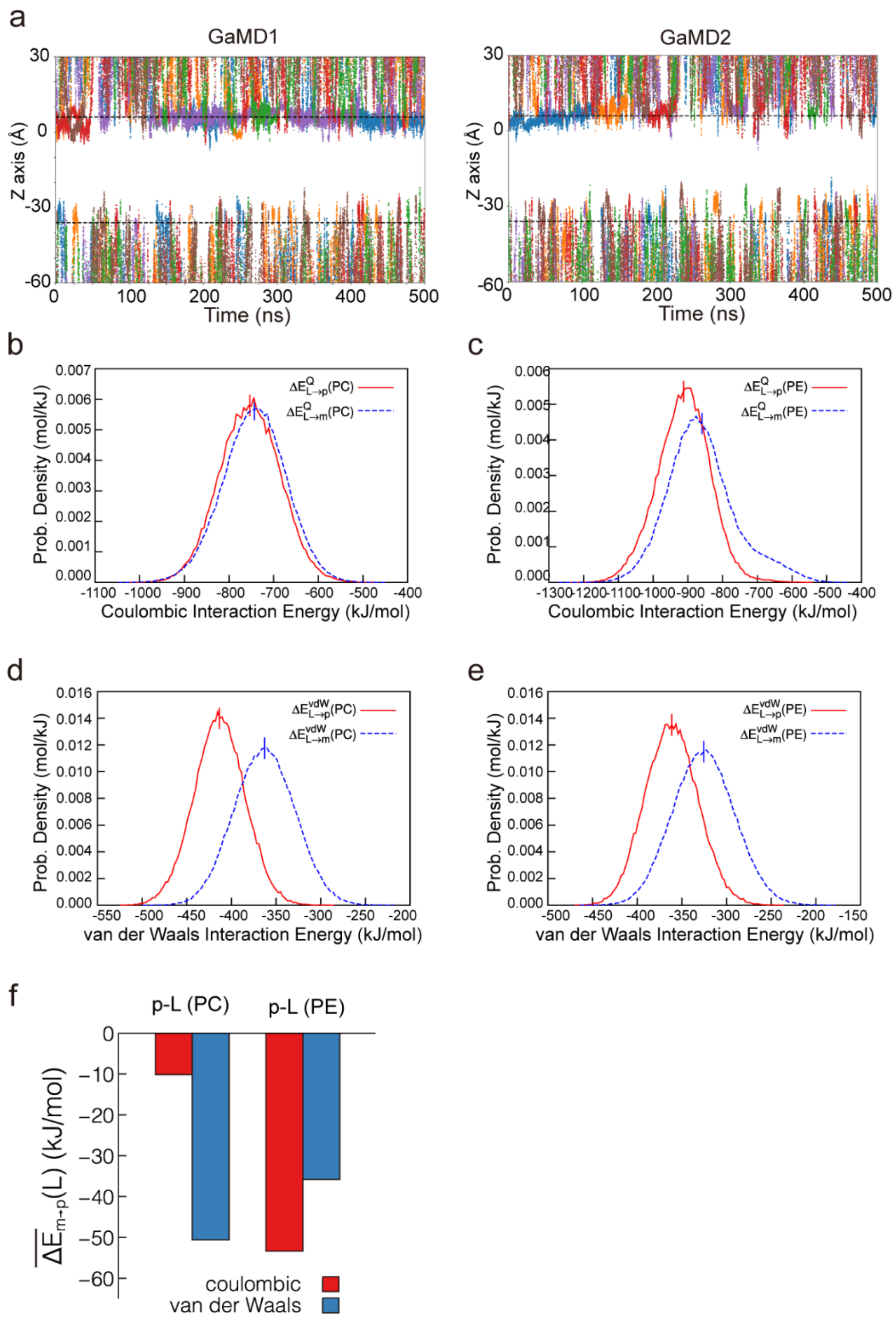


**Extended Data Fig.8 Molecular dynamics simulations of hTMEM87A.**

**a,** Representation of the Z coordinates of the $K^+$ atoms that bind in the negatively-charged luminal vestibule (NLV) along the GaMD simulations. The Z coordinates of the $K^+$ atoms from each binding event are indicated by different colored dots. The black dashed lines correspond to the upper and lower boundaries of the lipid bilayer membrane calculated as the averaged Z coordinates of the nitrogen atoms of the PE and PC molecules. Note that for the GaMD simulation 3 none binding event is explored, while for GaMD simulation 1 and 2 we explore 6 binding events each. When the binding takes place, the $K^+$ explores the NLV region (ca. 0-10 Z axis values) for a relatively long period of time.

**b-e**, The distributions of coulombic and van der Waals interaction energies for the PE/PC binding with hTMEM87A (solid lines) and for the PE/PC binding in a mixed lipid bilayer membrane (PC:PE=3:1) (dashed lines). The average interaction energies are marked by the vertical lines.

**f**, The comparison of average coulombic/van der Waals interaction energies of PE/PC transferring from membrane to protein. The transferring interaction energies are calculated as the differences between the average interaction energies in the **b-e** plots. The binding free energies in the main text can be calculated according to the LIE formula with 0.5 and 0.16 being the coefficients for the coulombic/van der Waals contributions, respectively.

**Extended Data Table 1. Cryo-EM data collection, refinement, and validation statistics**

| | #1 hTMEM87A (EMDB-34998) (PDB 8HSI) | #2 hTMEM87A-Gluc (EMDB-35017) (PDB 8HTT) |
|---|---|---|
| **Data collection and processing** | | |
| Magnification | 105,000x | 105,000x |
| Voltage (kV) | 300 | 300 |
| Electron exposure (e–/Å$^2$) | 67.72 | 68.1 |
| Defocus range (μm) | -0.8—1.9 | -0.8—1.9 |
| Pixel size (Å) | 0.849 | 0.849 |
| Symmetry imposed | C1 | C1 |
| Initial particle images (no.) | 8,101,104 | 6,035,205 |
| Final particle images (no.) | 445,198 | 201,915 |
| Map resolution (Å) | 3.14 | 3.6 |
| FSC threshold | 0.143 | 0.143 |
| Map resolution range (Å) | | |
| **Refinement** | | |
| Initial model used (PDB code) | - | - |
| Model resolution (Å) | | |
| FSC threshold | 0/0.143/0.5 | 0/0.143/0.5 |
| Model resolution range (Å) | 3.0/3.1/3.3 | 3.4/3.5/3.9 |
| Map sharpening *B* factor (Å$^2$) | 40.0 | 65.00 |
| Model composition | | |
| Non-hydrogen atoms | - | - |
| Protein residues | 405 | 389 |
| Ligands | BMA:1, NAG:6, CLR:2, L9Q:1 | BMA:1, NAG:6, GCO:1 |
| *B* factors (Å$^2$) | | |
| Protein | 122.92 | 163.62 |
| Ligand | 119.42 | 217.22 |
| R.m.s. deviations | | |
| Bond lengths (Å) | 0.028 | 0.003 |
| Bond angles (°) | 0.880 | 0.713 |
| Validation | | |
| MolProbity score | 2.05 | 2.11 |
| Clashscore | 14.83 | 18.66 |
| Poor rotamers (%) | 0.0 | 0.0 |
| Ramachandran plot | | |
| Favored (%) | 94.49 | 95.04 |
| Allowed (%) | 5.51 | 4.96 |
| Disallowed (%) | 0.0 | 0.0 |

**Extended Data Table 2. Constructs for recombinant protein expression**

| Construct | Expression vector | Information |
|---|---|---|
| hTMEM87A-WT-GFP-Strep | pcDNA3.4 TOPO | HindIII-hTMEM87A-WT (M1-E555)-TEV site-3C site-EGFP-)-thrombin site-Twin-Strep-tag-stop-XhoI |
| hTMEM87A-WT | pIRES2-DsRed | SalI-hTMEM87A-WT (M1-E555)-BamHI |
| hTMEM87A-Y237A | pIRES2-DsRed | SalI-hTMEM87A-Y237A (M1-E555)-BamHI |
| hTMEM87A-E272A | pIRES2-DsRed | SalI-hTMEM87A-E272A (M1-E555)-BamHI |
| hTMEM87A-K273A | pIRES2-DsRed | SalI-hTMEM87A-K273A (M1-E555)-BamHI |
| hTMEM87A-E279A | pIRES2-DsRed | SalI-hTMEM87A-E279A (M1-E555)-BamHI |
| hTMEM87A-E298A | pIRES2-DsRed | SalI-hTMEM87A-E298A (M1-E555)-BamHI |
| hTMEM87A-S301A | pIRES2-DsRed | SalI-hTMEM87A-S301A (M1-E555)-BamHI |
| hTMEM87A-K304A | pIRES2-DsRed | SalI-hTMEM87A-K304A (M1-E555)-BamHI |
| hTMEM87A-R305A | pIRES2-DsRed | SalI-hTMEM87A-R305A (M1-E555)-BamHI |
| hTMEM87A-R309A | pIRES2-DsRed | SalI-hTMEM87A-R309A (M1-E555)-BamHI |
| hTMEM87A-S415F | pIRES2-DsRed | SalI-hTMEM87A-S415F (M1-E555)-BamHI |
| hTMEM87A-S415W | pIRES2-DsRed | SalI-hTMEM87A-S415W (M1-E555)-BamHI |
| hTMEM87A-D442A | pIRES2-DsRed | SalI-hTMEM87A-D442A (M1-E555)-BamHI |

**Extended Data Table 3. Primer sequences used for cloning**

| Primer | Sequence (5'–3') |
|---|---|
| hTMEM87A-WT-GFP-strep-Foward | TTC TCT CCA CAG AAG CTT ATG GCG GCG GCT GCG TGG CT |
| hTMEM87A-WT-GFP-strep-Reverse | GTA CAG GTT TTC CTT AAG CTC CAT TTT GGA CCT TTC AAA GTG TGT GAT CAT TCG T |
| hTMEM87A-WT | TCG AAT TCT GCA GTC GAC ATG GCG GCG GCT GCG TGG CT |
| hTMEM87A-WT | AGG GAG AGG GGC GGA TCC CTC CAT TTT GGA CCT TTC AAA GTG TGT GAT CAT TCG T |
| hTMEM87A-Y237A-Forward | GTG ATG TGT ATT GTA GCT GTC CTG TTT GGT GTT CT |
| hTMEM87A-Y237A-Reverse | AGA ACA CCA AAC AGG ACA GCT ACA ATA CAC ATC AC |
| hTMEM87A-E272A-Forward | CTG GGA ATG CTT GAG AAA GCT GTC TTC |
| hTMEM87A-E272A-Reverse | GAA GAC AGC TTT CGC AAG CAT TCC CAG |
| hTMEM87A-K273A-Forward | GGA ATG CTT GAG AAA GCT GTC TTC TAT |
| hTMEM87A-K273A-Reverse | ATA GAA GAC AGC TGC CTC AAG CAT TCC |
| hTMEM87A-E279A-Forward | GTC TTC TAT GCG GCA TTT CAG AAT ATC CGA TAC |
| hTMEM87A-E279A-Reverse | GTA TCG GAT ATT CTG AAA TGC CGC ATA GAA GAC |
| hTMEM87A-E298A-Forward | TTG ATC CTT GCA GCG CTG CTT TCA GCA |
| hTMEM87A-E298A-Reverse | TGC TGA AAG CAG CGC TGC AAG GAT CAA |
| hTMEM87A-S301A-Forward | GCA GAG CTG CTT GCA GCA GTG AAA CGC |
| hTMEM87A-S301A-Reverse | GCG TTT CAC TGC TGC AAG CAG CTC TGC |
| hTMEM87A-K304A-Forward | CTT TCA GCA GTG AAA CGC TCA CTG GCT |
| hTMEM87A-K304A-Reverse | AGC CAG TGA GCG TGC CAC TGC TGA AAG |
| hTMEM87A-R305A-Forward | TCA GCA GTG AAA GCC TCA CTG GCT CGA |
| hTMEM87A-R305A-Reverse | TCG AGC CAG TGA GGC TTT CAC TGC TGA |
| hTMEM87A-R309A-Forward | CGC TCA CTG GCT GCA ACC CTG GTC ATC |
| hTMEM87A-R309A-Reverse | GAT GAC CAG GGT TGC AGC CAG TGA GCG |
| hTMEM87A-S415F-Forward | GCA GTG GCA GCA TTC ATT GTG TTT ATC ATC |
| hTMEM87A-S415F-Reverse | GAT GAT AAA CAC AAT GAA TGC TGC CAC TGC |

| | |
|---|---|
| hTMEM87A-S415W-Forward | GGC AGT GGC AGC ATA CAT TGT GTT TAT CAT C |
| hTMEM87A-S415W-Reverse | GAT GAT AAA CAC AAT GTA TGC TGC CAC TGC C |
| hTMEM87A-D442A-Forward | CTG TGG GTA GAC GAT GCC ATC TGG CGC |
| hTMEM87A-D442A-Reverse | GCG CCA GAT GGC AGC GTC TAC CCA CAG |
